# A histone variant condenses flowering plant sperm via chromatin phase separation

**DOI:** 10.1101/2021.09.14.460326

**Authors:** Toby Buttress, Shengbo He, Liang Wang, Shaoli Zhou, Lei Sun, Gerhard Saalbach, Martin Vickers, Pilong Li, Xiaoqi Feng

## Abstract

Sperm chromatin is typically transformed by protamines into a compact and transcriptionally inactive state. Flowering plant sperm cells lack protamines, yet have small, transcriptionally active nuclei with chromatin condensed by an unknown mechanism. Here we show that a histone variant, H2B.8, mediates sperm chromatin and nuclear condensation in *Arabidopsis thaliana*. Loss of H2B.8 causes enlarged sperm nuclei with dispersed chromatin, whereas ectopic expression in somatic cells produces smaller nuclei with aggregated chromatin, demonstrating that H2B.8 is sufficient for chromatin condensation. H2B.8 aggregates transcriptionally inactive AT-rich chromatin into phase-separated condensates, thus achieving nuclear compaction without reducing transcription. H2B.8 also intermixes inactive AT-rich chromatin and GC-rich pericentromeric heterochromatin, altering higher-order chromatin architecture. Altogether, our results reveal a novel mechanism of nuclear compaction via global aggregation of unexpressed chromatin. We propose that H2B.8 is a flowering plant evolutionary innovation that achieves nuclear condensation compatible with active transcription.

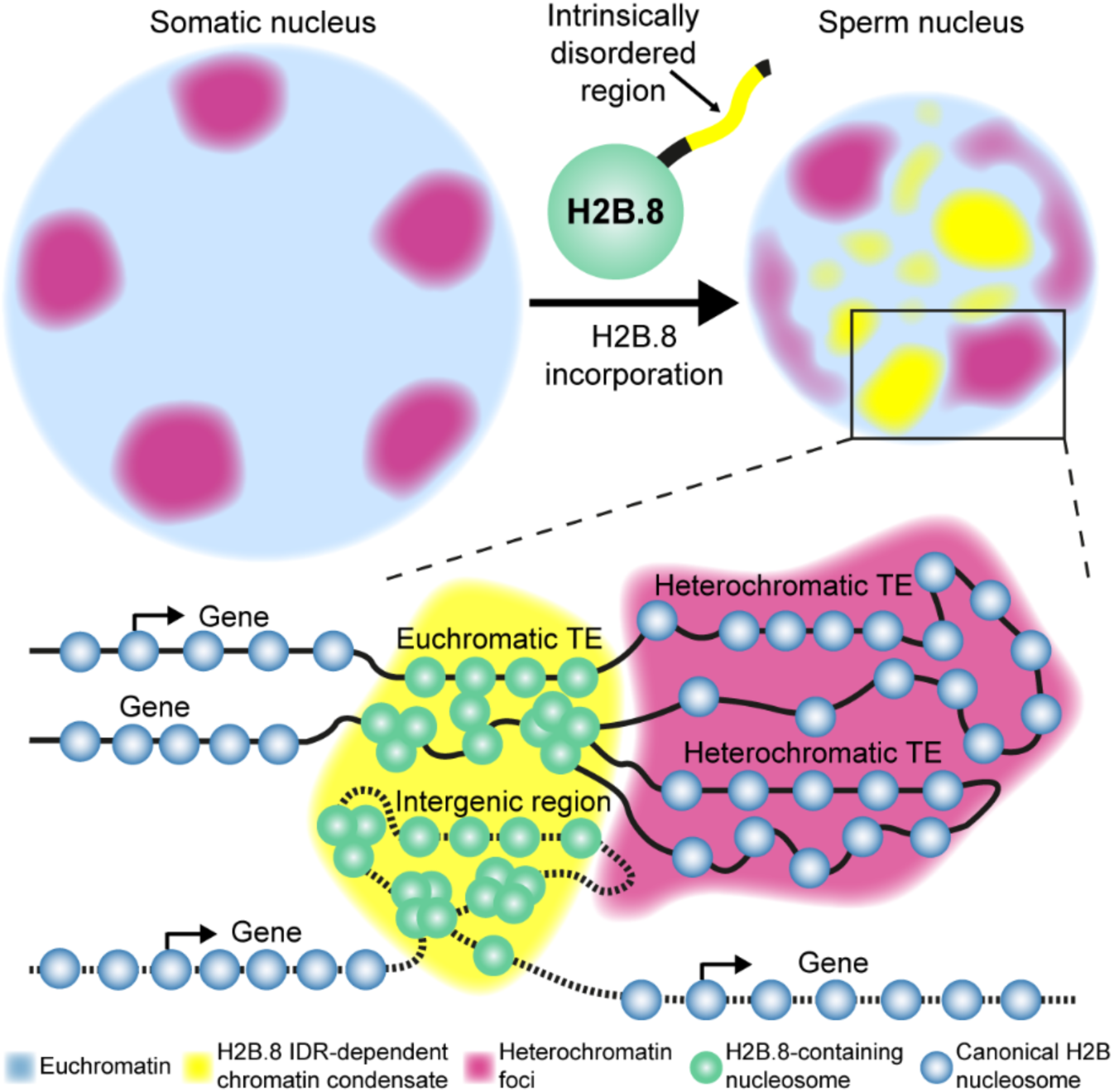

## MAIN TEXT

Sperm chromatin undergoes extensive condensation that is essential for male fertility in most animals. During animal sperm maturation, nearly all histones are replaced by small, arginine-rich protamines that achieve a denser packaging of the DNA (*1, 2*). In mammalian sperm for example, histones are retained at only 5-15% of the genome, whereas the majority of regulatory information carried by histones is lost (*1, 3*). Unwinding DNA from histones requires DNA strand breaks, which are observed in great quantities during this process (*4, 5*). Furthermore, protamines preclude transcription (*6, 7*). The extreme chromatin compaction enabled by protamines protects genome integrity from genotoxic factors and achieves a small and hydrodynamic sperm head that enhances swimming ability (*2, 7*). The adoption of such a drastic process illustrates the high evolutionary pressure on sperm fitness (*8–10*).

Sperm condensation also occurs in another large group of multicellular eukaryotes, plants. Similar to animals, green algae and non-seed plants such as liverworts, mosses, and ferns, produce motile sperm, which swim through water to reach the egg cell (*11*). Consistent with the theory that protamine-mediated sperm condensation evolved to facilitate swimming (*2*), sperm nuclei in these species are highly condensed by protamines and protamine-like proteins and transcribe very little RNA, if any (*12–16*).

Diverged from other land plant species approximately 150 million years ago, flowering plants no longer rely on water for fertilization (*11*). Flowering plants produce immotile, transcriptionally active sperm (*17, 18*). The sperm cells are encapsulated in pollen grains, which are produced by mitotic divisions of the haploid meiotic product called the microspore (*11*). The microspore divides once to produce a vegetative cell and a generative cell, the latter of which subsequently divides to generate two sperm cells (*11*). During fertilization, the vegetative cell develops into a pollen tube that delivers the sperm to the egg apparatus (*19*). Consistent with the high metabolic activity required for powering pollen tube growth, the vegetative cell chromatin is highly decondensed and transcriptionally active (*20–23*). In contrast, sperm has highly condensed, histone-based chromatin and small nuclei (*24, 25*). In the absence of protamines, the mechanism of sperm chromatin compaction in flowering plants is unknown.

To understand the mechanism underlying sperm condensation in flowering plants, we performed super-resolution imaging and comparative proteomics on *Arabidopsis thaliana* sperm, vegetative and somatic cells. Through this, we identified a specifically expressed histone variant, H2B.8, that colocalizes with chromatin aggregates in the sperm nucleoplasm. *h2b.8* mutant sperm nuclei are enlarged with decondensed chromatin, while the ectopic expression of H2B.8 in somatic cells causes the opposite phenotype, demonstrating that H2B.8 is required and sufficient for nuclear and chromatin compaction. H2B.8 compacts chromatin via a phase separation mechanism that is dependent on a conserved intrinsically disordered region (IDR). H2B.8 specifically concentrates unexpressed AT-rich chromatin, thereby reducing nuclear volume while maintaining transcription. Collectively, our results elucidate flowering plant sperm condensation and reveal a novel mechanism of nuclear compaction via phase-separation-mediated chromatin aggregation.

### Chromatin aggregates are observed throughout sperm nucleoplasm

Previously, *Arabidopsis* sperm chromatin was shown to be highly condensed based on strong DAPI staining and small nuclear size, compared with somatic and vegetative cells (*24*). To characterize sperm chromatin in more detail, we examined *Arabidopsis* sperm nuclei using super-resolution 3D Structured Illumination Microscopy (3D-SIM). Distinct chromatin aggregates were observed throughout the nucleoplasm in sperm (Fig. 1A). In contrast, the vegetative chromatin is more homogenous, and so is the leaf cell chromatin except for the condensed heterochromatin foci at the nuclear periphery (Fig. 1A).

**Fig. 1.**
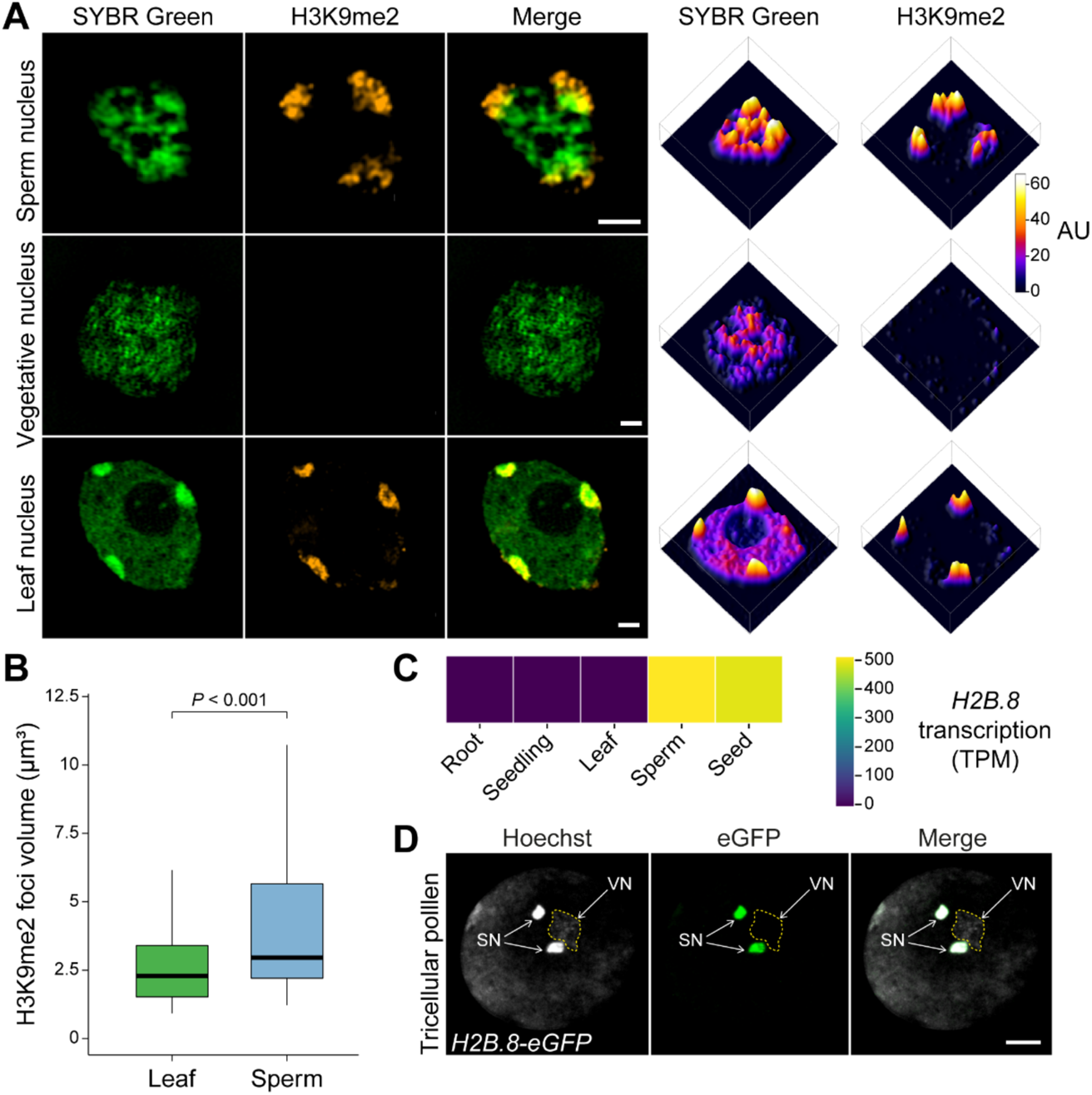
Sperm chromatin is aggregated and contains a specific histone variant H2B.8. (**A**) Super-resolution 3D-SIM images and associated intensity profiles of wild-type sperm and vegetative nuclei from pollen and a leaf nucleus. DNA is stained with SYBR Green (green) and H3K9me2 is immunolocalized (orange). AU, arbitrary units. Scale bars, 1 μm. (**B**) Volumes of H3K9me2-enriched heterochromatin foci in leaf and sperm nuclei. *P*-value, independent two-sample t-test; n = 30 nuclei each. (**C**) *H2B.8* transcription in indicated tissues and cells. TPM, transcripts per million. (**D**) Confocal images of *pH2B.8::H2B.8-eGFP* pollen, in which eGFP signal is specific to the sperm nuclei (SN). VN, vegetative nucleus (outlined in a dashed line). Scale bar, 5 μm.

To understand the composition of chromatin within sperm aggregates, we performed immunostaining against histone H3 lysine 9 dimethylation (H3K9me2), a modification associated with silenced heterochromatin (*26*). We found some of the larger aggregates situated at the sperm nuclear periphery colocalize with H3K9me2 signal (Fig. 1A). This shows that heterochromatin domains persist in the sperm, as previously reported (*22*). However, these heterochromatin foci are enlarged in the sperm in comparison to leaf cells (Figs. 1, A and B), despite sperm, being haploid, having only half as much DNA as leaf cells. Such enlargement indicates a reduced level of heterochromatin condensation in sperm. Besides the heterochromatin foci, the rest of the chromatin aggregates in sperm are H3K9me2 depleted (Fig. 1A), suggesting the involvement of a novel mechanism that compacts the less heterochromatic part of sperm chromatin.

### Canonical H2B is partially replaced by the H2B.8 variant in sperm

To investigate the mechanism of sperm chromatin compaction, we searched for sperm-specific chromatin factors by performing mass spectrometry on leaf nuclei and FACS-isolated sperm and vegetative nuclei. We identified a variant of histone H2B, H2B.8 (encoded by *AT1G08170*), which constitutes 12.6% of H2Bs in sperm but is absent in vegetative or leaf nuclei (figS. S1A). Consistently, RNA-seq experiments detected abundant *H2B.8* transcript in the sperm, but none from somatic tissues such as leaves, roots and whole seedlings (Fig. 1C). To further examine the protein expression pattern during development, we generated a reporter line by expressing H2B.8-eGFP fusion with the native *H2B.8* promoter in *Arabidopsis* (*pH2B.8::H2B.8-eGFP*). Confocal imaging shows that H2B.8 is incorporated into sperm following the second pollen mitotic division, when nuclei are compacted and chromatin aggregates (Fig. 1D and figS. S1B). H2B.8 is rapidly lost after fertilization but reappears during seed maturation (figS. S1C). No H2B.8-eGFP was observed in any other cell or tissue except seeds (figS. S1, B and C), consistent with recently published analyses of H2B expression (*27, 28*).

### H2B.8 drives chromatin aggregation and nuclear condensation

Because the presence of H2B.8 correlates with nuclear and chromatin condensation, we hypothesized that H2B.8-induced chromatin compaction is responsible for sperm condensation. To test this hypothesis, we generated two independent *h2b.8* CRISPR knockout mutants (*h2b.8-1* and *h2b.8-2*; figS. S2A) and examined their sperm phenotype via confocal and 3D-SIM imaging. We found that sperm nuclei of the two *h2b.8* mutants are about 40% larger than those of the wild type (Fig. 2A). Additionally, in support of our hypothesis, chromatin is more homogenous with reduced aggregation in *h2b.8* mutants (Fig. 2B and figS. S2B). To pinpoint the role of *h2b.8* mutations in causing these phenotypes, we expressed *pH2B.8::H2B.8-Myc* and *pH2B.8::H2B.8-eGFP* transgenes in the *h2b.8* mutant (*h2b.8-1*, unless specified all *h2b.8* mutants hereafter refer to this allele). Both transgenes successfully rescued the sizes of *h2b.8* sperm nuclei to wild-type levels (Fig. 2A), confirming that H2B.8 drives nuclear condensation. Chromatin aggregates are also restored in *pH2B.8::H2B.8-eGFP h2b.8* sperm (Fig. 2C). Furthermore, these restored aggregates colocalize with H2B.8-eGFP (Fig. 2C), suggesting that H2B.8 is directly involved in forming the aggregates.

**Fig. 2.**
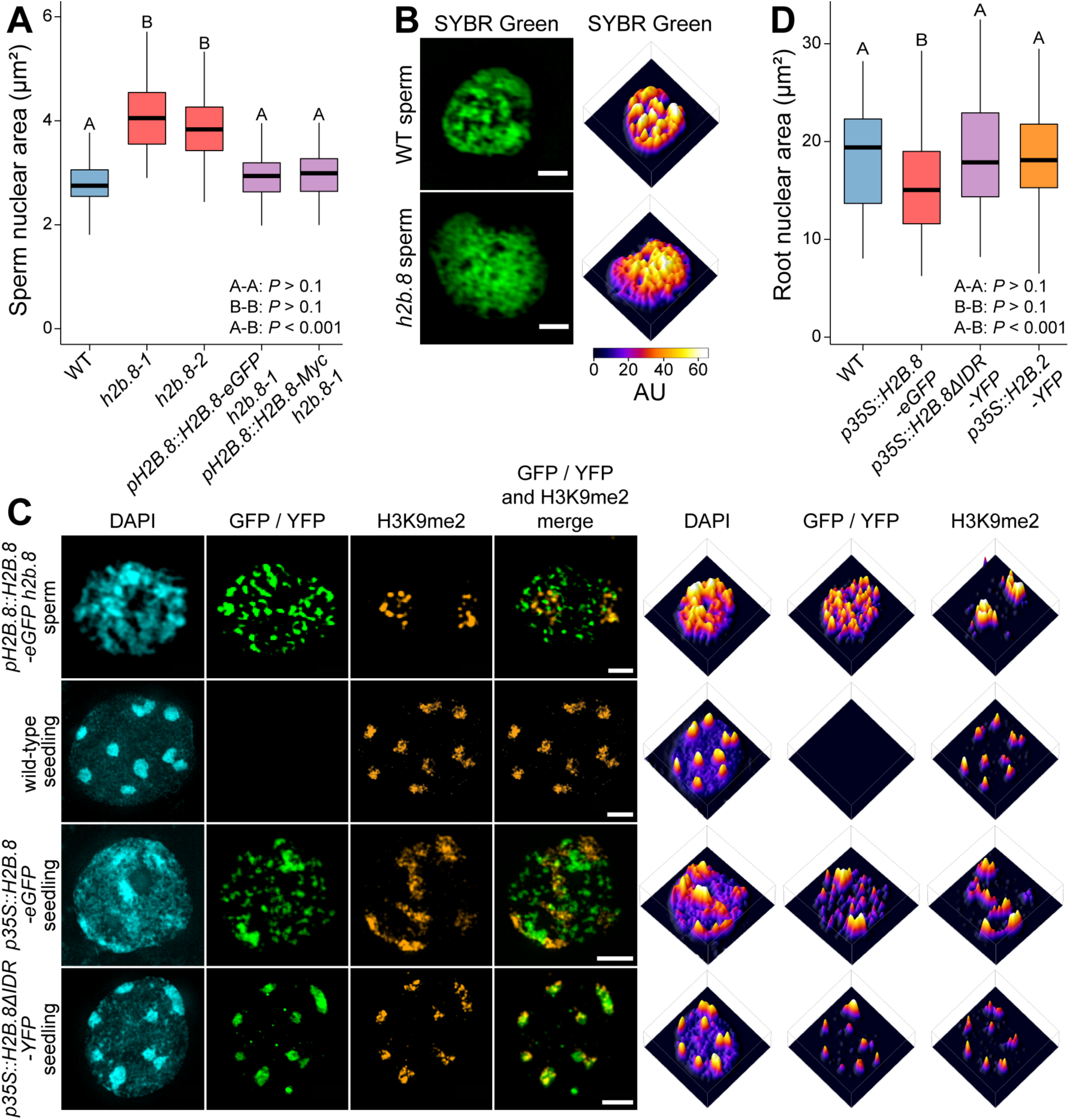
H2B.8 is required and sufficient to drive chromatin and nuclear compaction. (**A**) Sperm nuclear sizes in wild type (WT), two independent *h2b.8* CRISPR mutants and complementation lines of the *h2b.8-1* mutant. *P*-values, ANOVA followed by individual two-sample Tukey tests. Boxplots marked as A and B are significantly different between groups (*P* < 0.001) but not within the group (*P* > 0.1). n = 80 (WT, *h2b.8-1*, *pH2B.8::H2B.8-Myc h2b.8-1*), 77 (*h2b.8-2*), and 79 (*pH2B.8::H2B.8-eGFP h2b.8-1*). (**B** and **C**) Super-resolution 3D-SIM images and associated intensity profiles of WT and *h2b.8* (all *h2b.8* refers to *h2b.8-1* unless specified otherwise) sperm nuclei (B), and sperm and seedling nuclei of indicated genotypes (C). AU, arbitrary units. Scale bars, 1 μm (B), 1 μm (C, upper panels) and 2 μm (C, middle and lower panels). (**D**) Root nuclear sizes in WT and lines ectopically expressing H2B.8, H2B.8ΔIDR and H2B.2. *P*-values, ANOVA followed by individual two-sample Tukey tests. Boxplots marked as A and B are significantly different between groups (*P* < 0.001) but not within the group (*P* > 0.1). n = 81 (WT), 96 (*p35S::H2B.8-eGFP*), 156 (*p35S::H2B.8ΔIDR-YFP*), and 129 (*p35S::H2B.2-YFP*).

To test if H2B.8 is sufficient to drive chromatin aggregation and nuclear compaction, we ectopically expressed H2B.8 using a strong constitutive promoter (*p35S*). Distinctive chromatin aggregates that co-localize with H2B.8-eGFP are induced in *p35S::H2B.8-eGFP* seedling cell nuclei (Fig. 2C). Furthermore, *p35S::H2B.8-eGFP* expression reduced nuclear size in root cells by 22.4% (Fig. 2D). These results demonstrate that H2B.8 is sufficient for chromatin and nuclear condensation.

### H2B.8 aggregates chromatin via IDR-dependent phase separation

H2B.8 is distinguished from other *Arabidopsis* H2B variants by a much longer N-terminal tail that contains a 93 amino acid intrinsically disordered region (IDR; figS. S3, A and B). Phylogenetic analysis revealed that H2B.8 is specific to flowering plants and is present in all flowering plant species with published genomes except the most basal *Amborella trichopoda* (Fig. 3A; Table S1). Notably, all identified H2B.8 homologs share the insertion of an IDR in the histone tail (Fig. 3A; Table S1), suggesting its functional importance.

**Fig. 3.**
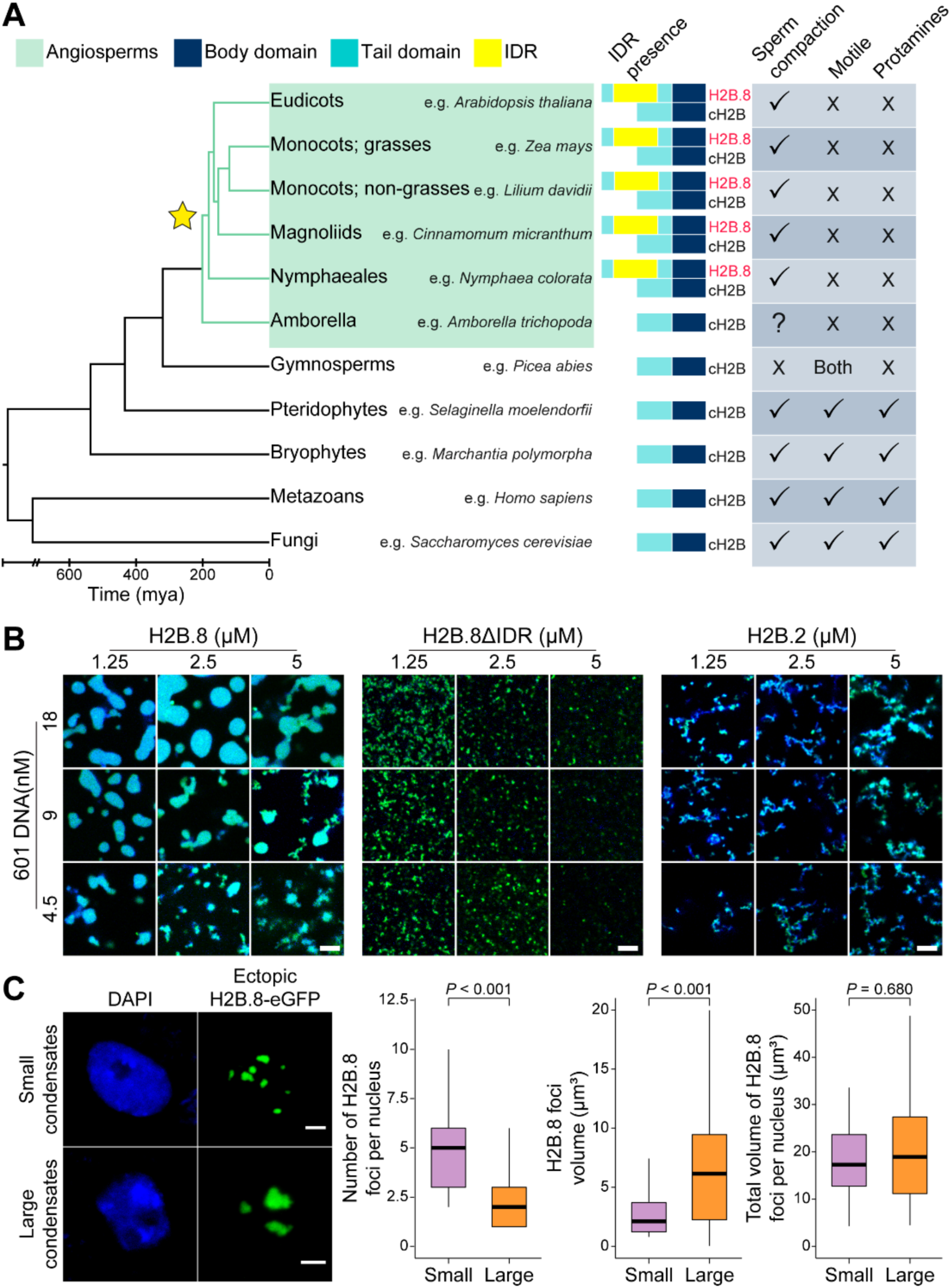
H2B.8 condenses chromatin via IDR-dependent phase separation. (**A**) Phylogenetic tree illustrating H2B.8 evolution (marked by a star). Sperm chromatin compaction state, sperm motility and the presence of protamines or protamine-like proteins are denoted for represented eukaryote lineages. (**B**) *In vitro* phase separation assays of purified histone H2B.8, H2B.8ΔIDR and H2B.2 (Alexa Fluor 488; green) with Widom 601 DNA (DAPI; blue) under physiological salt conditions. Scale bars, 5 μm. (**C**) Confocal images and quantification of H2B.8 condensates in *p35S::H2B.8-eGFP* seedlings. *P*-value, independent two-sample t-test. n = 30 and 71 for nuclei with small and large H2B.8 condensates, respectively. Scale bars, 2 μm.

IDRs have been demonstrated to drive the formation of biomolecular condensates by phase separation (*29–33*). Moreover, phase separation of IDR-containing proteins has been shown to drive the formation and condensation of heterochromatin foci (*34–37*). Based on this knowledge and the distinctive H2B.8 foci in sperm and seedling cells (Fig. 2C), we hypothesized that H2B.8 aggregates chromatin via IDR-mediated phase separation.

Consistent with this hypothesis, we observed that H2B.8 can form homogenous condensates *in vitro* at low concentrations in a DNA-dependent manner (Fig. 3B). The IDR is critical for condensate formation as H2B.8 without the IDR (H2B.8ΔIDR) and a canonical H2B (H2B.2) fail to undergo phase separation *in vitro* under physiological salt conditions (Fig. 3B). To investigate the physical nature of H2B.8 condensates, we performed fluorescence recovery after photobleaching (FRAP). H2B.8 condensates fail to recover after photobleaching both *in vitro* and *in vivo* (figS. S3, C and D), indicating they are gel-like in nature. Consistent with such property, H2B.8 droplets fuse slowly *in vitro* (figS. S3E).

We next examined the ectopically induced H2B.8 condensates in *p35S::H2B.8-eGFP* root cell nuclei. These condensates form a gradient of sizes and numbers among nuclei. Approximately 30% of nuclei contain numerous small H2B.8 condensates; the remaining 70% have fewer and larger condensates (Figs. 2C and 3C). The total volume of H2B.8 condensates is comparable between the two types of nuclei (Fig. 3C), suggesting that H2B.8-containing chromatin condensates are fusing over time.

To test whether H2B.8 phase separation is required for chromatin condensation *in vivo*, we ectopically expressed H2B.8ΔIDR (*p35S::H2B.8ΔIDR-YFP*). In contrast to the effect of full-length H2B.8, H2B.8ΔIDR expression did not induce chromatin aggregation and had no effect on nuclear size in root cells (Figs. 2, C and D), indicating that chromatin and nuclear condensation are dependent on phase separation of H2B.8.

### H2B.8 is located within AT-rich, transcriptionally inactive sequences

To further understand H2B.8 activity, we determined H2B.8 genomic localization via native ChIP-seq on *pH2B.8::H2B.8-eGFP h2b.8* pollen using GFP antibodies. This identified H2B.8 peaks occupying roughly 17% of the sperm genome, comparable with our mass spectrometry results that approximately 13% of canonical H2B is replaced by H2B.8 in sperm (figS. S1A). H2B.8 is most enriched within so-called euchromatic transposable elements (TEs; Figs. 4, A to C), which are AT-rich and depleted of H3K9me2 and other heterochromatic marks (*38*). Heterochromatic TEs that are typically GC- and H3K9me2-rich (*38–40*) have comparatively little H2B.8 (Figs. 4, A to C). H2B.8 is excluded from the bodies of transcribed genes compared to inactive genes and intergenic regions, and H2B.8 enrichment and gene transcription are anticorrelated (Figs. 4, A to D). H2B.8 distribution along chromosomes follows that of euchromatic TEs, with H2B.8 most abundant at the edges of pericentromeric regions (Fig. 4E and figS. S4A).

**Fig. 4.**
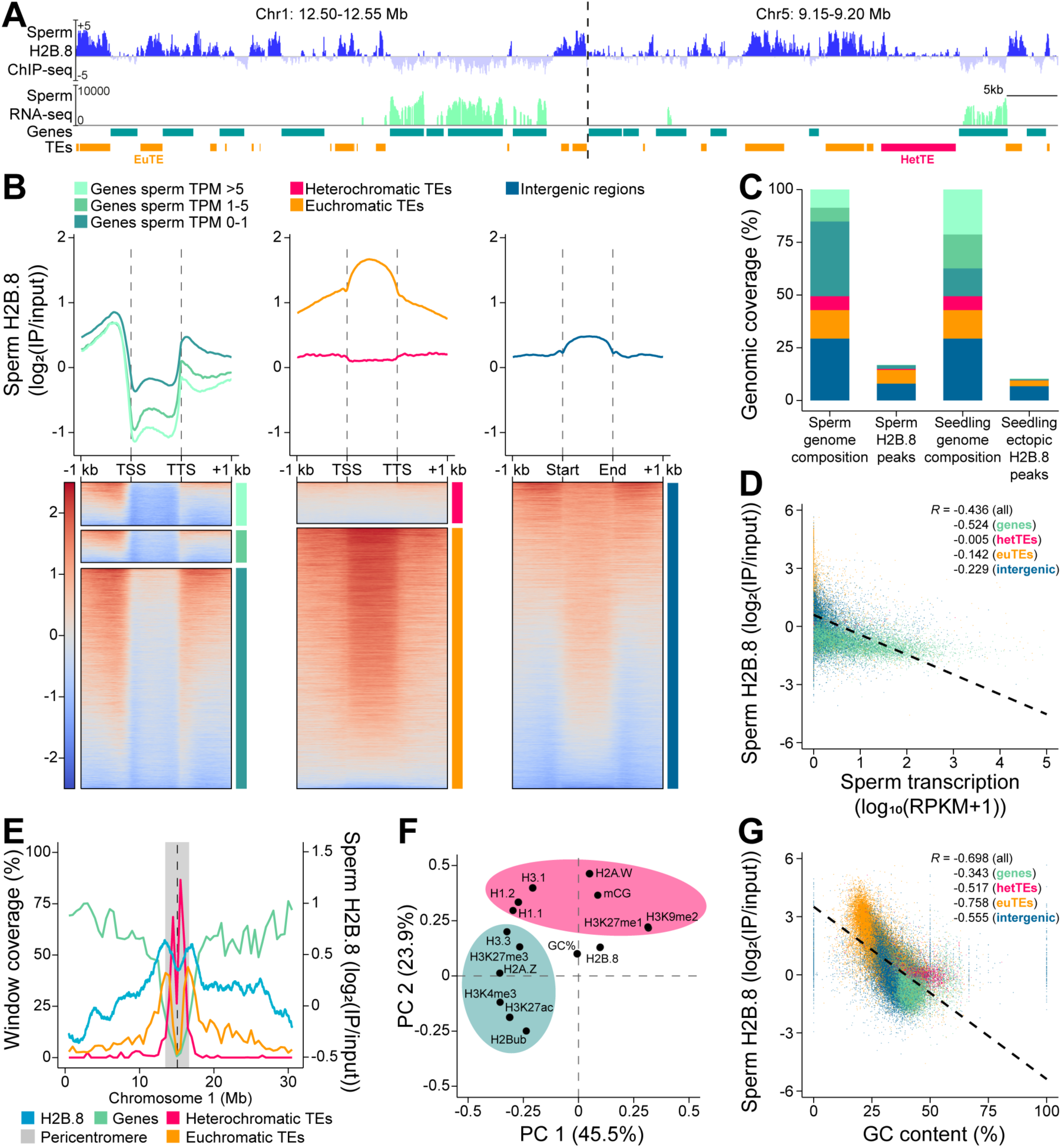
H2B.8 is localized in AT-rich TEs and transcriptionally inactive intergenic regions. (**A**) Genome snapshots of H2B.8 abundance in sperm (log_2_(IP/input)), sperm cell transcription (log_2_(RPKM)), and gene and TE annotations (orange, euchromatic TE; magenta, heterochromatic TE) over representative 50 kb regions. RPKM, Reads Per Kilobase of transcript per Million mapped reads. (**B**) Profiles and associated heatmaps of sperm H2B.8 enrichment over genes (grouped by sperm cell expression), TEs (grouped by chromatin state) and intergenic regions. (**C**) Proportions (%) of the genome covered by genes, TEs and intergenic regions in wild-type sperm and seedling, and by respective H2B.8 peaks in *pH2B.8::H2B.8-eGFP h2b.8* sperm and *p35S::H2B.8-eGFP* seedling. Same color coding is used as in (B). (**D** and **G**) Scatterplots showing anticorrelation of sperm H2B.8 enrichment with sperm transcription (D) or GC content (G) among indicated genomic features. *R*, Spearman’s Rank. (**E**) Coverage of genes, euchromatic TEs and heterochromatic TEs (left Y axis; 500 kb windows) and H2B.8 enrichment in sperm (right Y axis; 1 kb windows) along Chromosome 1. Chromosomes 2-5 are shown in Figure S4A. (**F**) Principal Component Analysis of H2B.8 abundance in *p35S::H2B.8-eGFP* seedlings with other chromatin marks. The green and pink shaded areas represent euchromatic and heterochromatic marks, respectively.

To further understand the chromatin preferences of H2B.8 and whether its localization pattern is intrinsically determined by H2B.8 or other sperm-specific components, we performed native ChIP-seq with seedlings of the ectopic H2B.8 expression line (*p35S::H2B.8-eGFP*). This revealed an analogous H2B.8 localization pattern to that in sperm, with enrichment in euchromatic TEs and intergenic regions (Fig. 4C and figS. S4, B and C). Together with the ability of ectopic H2B.8 to condense root cell chromatin and nuclei (Figs. 2, C and D), this suggests that H2B.8 deposition is not reliant on sperm-specific factor(s). Next, utilizing the available seedling epigenomic data, we explored H2B.8 associations with other chromatin features. Principal component analysis (PCA) revealed that H2B.8 clusters with neither permissive nor repressive chromatin modifications (Fig. 4F). The best predictor of H2B.8 localization is GC content, with which a strong anti-correlation exists (Figs. 4, F and G and figS. S4D). This likely explains why genes and H3K9me2-enriched heterochromatic TEs, both GC-rich, are generally depleted of H2B.8 (Figs. 4, B and G). Among the GC-poor elements, mostly euchromatic TEs and intergenic regions, H2B.8 anti-correlates with transcription and associated histone marks (e.g. H3K4me3; Fig. 4D and figS. S4, E and F). In all, our results suggest that H2B.8 localization is mostly driven by GC content and transcription rather than sperm-specific factors.

### H2B.8 condenses chromatin without affecting transcription

Chromatin condensation is frequently associated with transcriptional repression (*41–43*). However, the localization of H2B.8 in the non-transcribing parts of the genome suggests that it may not adversely affect gene expression. To test this hypothesis, we isolated wild-type and *h2b.8* mutant sperm cells and performed RNA sequencing. Among the 12198 genes expressed in either wild-type or *h2b.8* sperm, none have significantly altered expression in *h2b.8* (Fig. 5A). Likewise, we did not find any TE significantly misregulated in *h2b.8* sperm (Fig. 5B). RNA sequencing of wild-type and *p35S::H2B.8-eGFP* seedlings further supported the negligible effect of H2B.8 on transcription (figS. S5, A and B). Therefore, unlike protamines, which condense animal sperm at the expense of transcriptional potential (*6, 7*), H2B.8 condenses plant sperm without suppressing transcription.

**Fig. 5.**
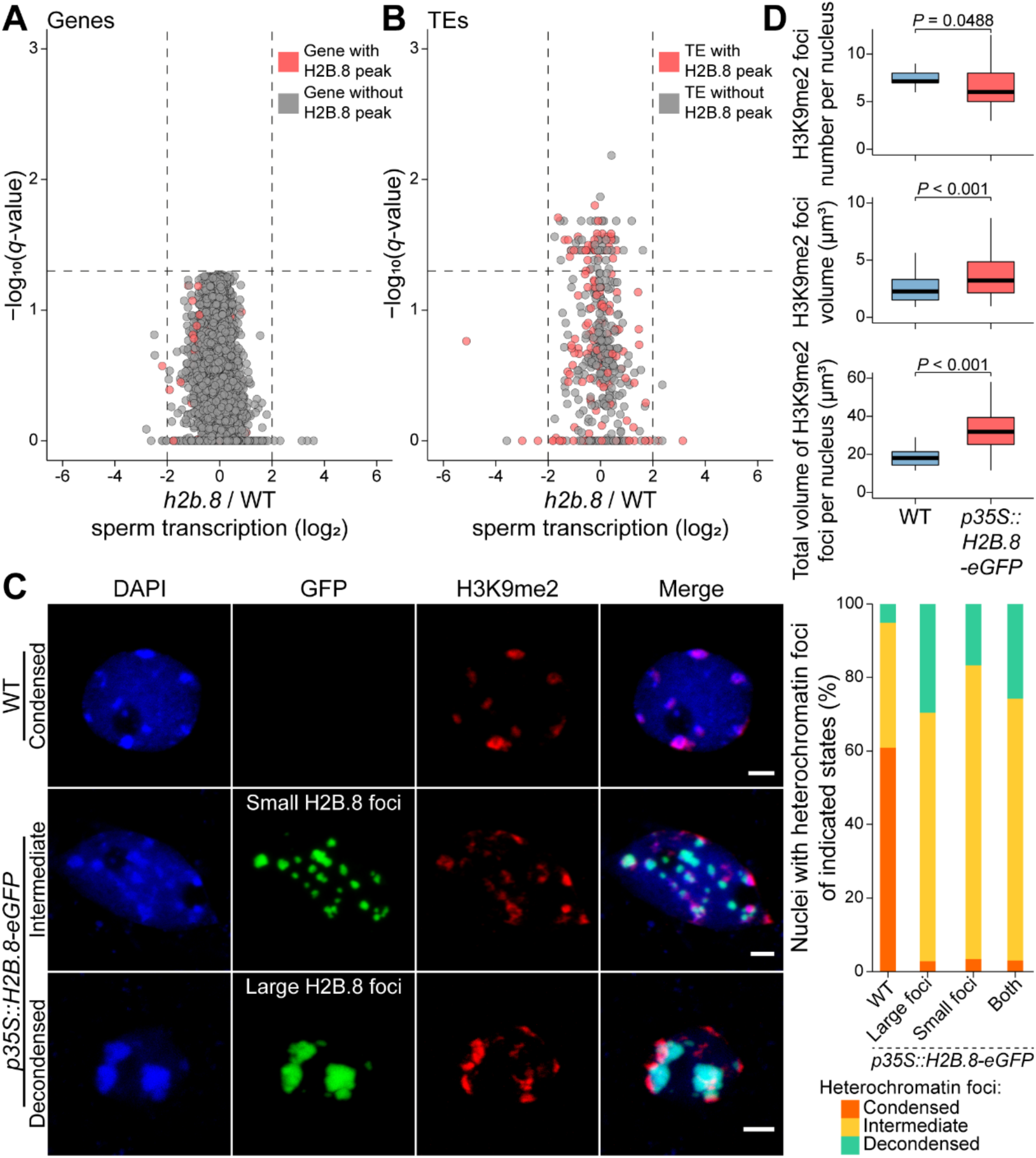
H2B.8-induced chromatin aggregation does not suppress transcription but decondenses heterochromatin foci. (**A** and **B**) Volcano plots showing differential gene (A) and TE (B) expression between *h2b.8* mutant and wild-type (WT) sperm cell. Differentially expressed genes/TEs were defined as log_2_(*h2b.8*/WT TPM fold change) ≥2 or ≤-2 and *q* < 0.05 (likelihood-ratio test). n = 12198 (A) and 480 (B). TPM, transcripts per million. (**C**) Confocal images and quantification of seedling nuclei of indicated genotypes with condensed, intermediately condensed or decondensed heterochromatin foci (measured by H3K9me2 signal). n = 138 (WT), and 71, 30 and 101 (*p35S::H2B.8-eGFP* nuclei with large, small, and both large and small H2B.8 foci, respectively). (**D**) Quantification of H3K9me2-enriched heterochromatin foci in WT and *p35S::H2B.8-eGFP* seedling nuclei. *P*-value, independent two-sample t-test. n = 138 and 101 for WT and *p35S::H2B.8-eGFP*, respectively.

### H2B.8 phase separation decondenses heterochromatin

Besides the overall chromatin compaction in sperm, we observed slight decondensation of heterochromatin foci via 3D-SIM (Figs. 1, A and B). To understand if this is caused by H2B.8, we performed immunostaining of *p35S::H2B.8-eGFP* seedling nuclei using H3K9me2 antibodies. Consistent with H2B.8 ChIP-seq data (figS. S4, B and C) and our observations in *pH2B.8::H2B.8-eGFP h2b.8* sperm (Fig. 2C), H2B.8 condensates are largely devoid of H3K9me2 and distinct from heterochromatin foci (Fig. 5C). However, the heterochromatin foci are enlarged and decondensed in *p35S::H2B.8-eGFP* seedling nuclei in comparison to the wild type (Figs. 5, C and D). Although most wild-type nuclei show highly condensed heterochromatin foci, these are rarely found in *p35S::H2B.8-eGFP* seedling nuclei (Fig. 5C). The majority of *p35S::H2B.8-eGFP* seedling nuclei exhibit moderately dispersed heterochromatin foci (Figs. 5, C and D), reminiscent of those in sperm (Figs. 1, A and B). This suggests that H2B.8 causes heterochromatin foci to decondense.

We further found that decondensation of heterochromatin foci is dependent on H2B.8 phase separation, as the expression of H2B.8ΔIDR does not affect heterochromatin (Fig. 2C). As heterochromatin foci are themselves phase-separated condensates (*35, 36, 44, 45*), this suggests interactions between the two types of condensates. Indeed, although H2B.8 and heterochromatic condensates are mostly distinct, some physical associations are observed (Fig. 5C). Collectively, our results show that condensation of chromatin via H2B.8 phase separation affects heterochromatin condensation, likely because H2B.8-associated AT-rich euchromatic TEs are interspersed with heterochromatic TEs in pericentromeric regions (Fig. 4E and figS. S4A).

### H2B.8 affects intra- and inter-chromosomal interactions

To understand how H2B.8 mediates chromatin compaction, we performed genome-wide chromosome conformation capture analysis (Hi-C) (*46*) on seedlings ectopically expressing H2B.8 (*p35S::H2B.8-eGFP*) and wild-type controls. The Hi-C libraries were sequenced to single kilobase resolution (figS. S6A), and our wild-type contact matrices are comparable to previously published experiments (figS. S6B) (*47, 48*). As shown before (*49–51*), topologically associating domains are absent in *Arabidopsis*, but telomeres are frequently associated, as are centromeres (figS. S6C).

The comparison of *p35S::H2B.8-eGFP* and wild-type Hi-C data revealed alterations of higher-order chromatin architecture. Within chromosomes, ectopic H2B.8 caused increased short-range contacts (200 kb – 1.1 Mb) and depleted long-range contacts (> 1.1 Mb) (figS. S6D). Short-range intrachromosomal interactions are indicative of chromatin compaction, suggesting that H2B.8 principally forms aggregates by concentrating linearly proximal regions (Fig. 6A and figS. S6D). In support of this, the increase of short-range contacts strongly correlates with local H2B.8 abundance (Spearman’s *R* = 0.97; Fig. 6B).

**Fig. 6.**
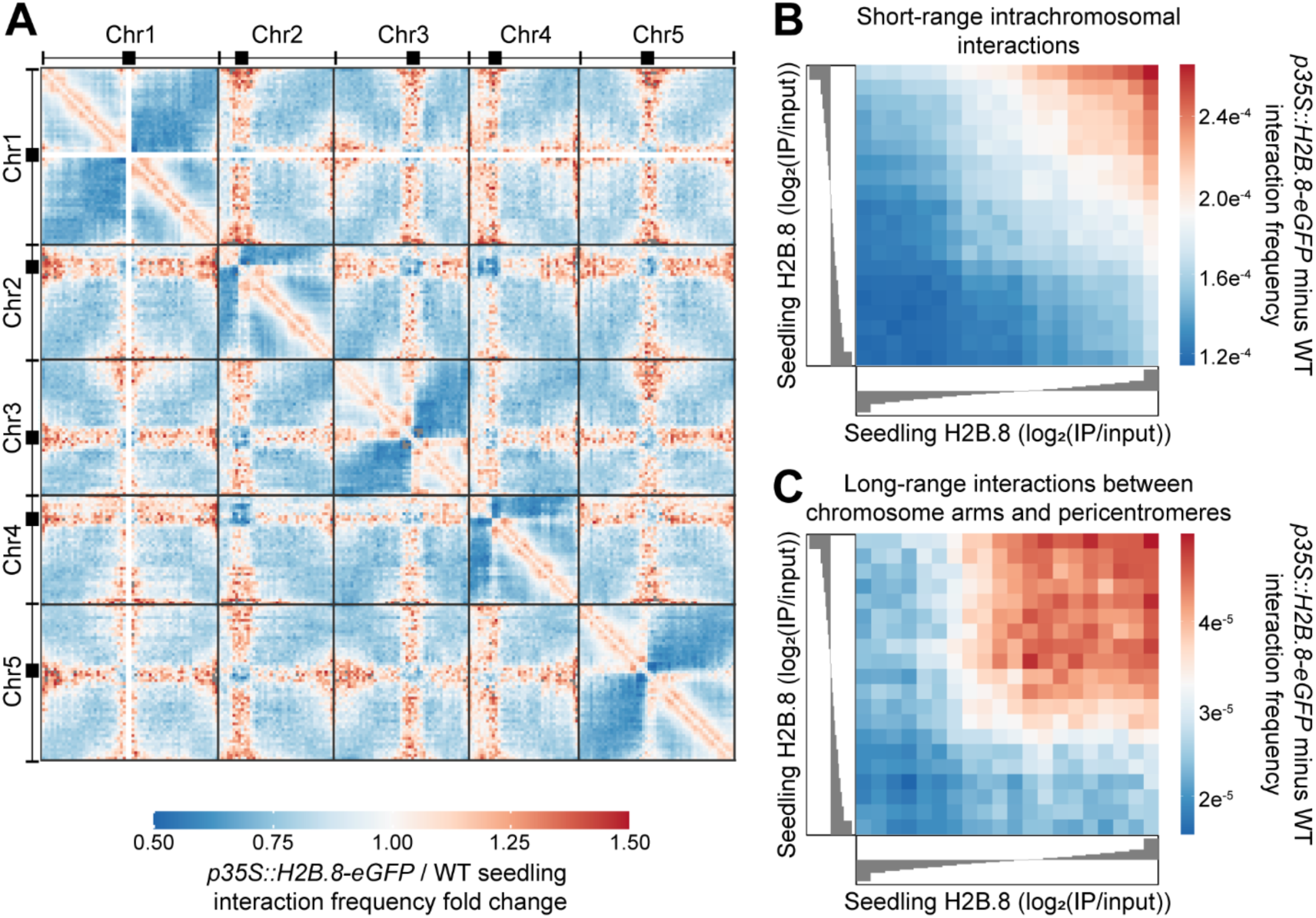
H2B.8 drives aggregation of euchromatic arms and pericentromeric heterochromatin. (**A**) Genome-wide interaction frequency fold change heatmap between wild-type (WT) and *p35S::H2B.8-eGFP* seedlings at 500 kb resolution. (**B**) Short-range intrachromosomal interaction frequency difference between *p35S::H2B.8-eGFP* and WT over quantiles of seedling H2B.8 enrichment (log_2_(IP/input)). Spearman’s *R* = 0.974. (**C**) Long-range interaction frequency difference between *p35S::H2B.8-eGFP* and WT between chromosome arms and pericentromeric regions over quantiles of seedling H2B.8 enrichment (log_2_(IP/input)). Spearman’s *R* = 0.890.

Contacts between pericentromeric regions (heterochromatin) and distal chromosomal arms (euchromatin) are also increased in *p35S::H2B.8-eGFP* seedlings (Fig. 6A). Concomitantly, interchromosomal interactions between pericentromeres are reduced (Fig. 6A). These alterations are consistent with the cytologically observed heterochromatin decondensation and association of heterochromatin foci and H2B.8 condensates (Figs. 2C and 5C). The effect of H2B.8 is local, as the interactions between pericentromeric regions and chromosomal arms increase at regions with abundant H2B.8 (Fig. 6C). These observations support the hypothesis that dispersal of heterochromatin foci is caused by H2B.8-mediated aggregation of euchromatic TEs that are abundant in and near pericentromeric regions (Fig. 4E and figS. S4A). Taken together, our Hi-C and ChIP-seq data demonstrate that H2B.8 achieves a form of global chromatin compaction via binding and aggregating transcriptionally inactive AT-rich sequences dispersed throughout the genome.

## Discussion

Our results reveal a mechanism of chromatin compaction driven by H2B.8-induced phase separation (figS. S7). Chromatin aggregates in the sperm nucleus are diminished in *h2b.8* mutants (Fig. 2B and figS. S2B), whereas ectopically expressed H2B.8 in somatic cells is sufficient to induce chromatin aggregates in an IDR-dependent manner (Fig. 2C). Hi-C and cytological observations reveal that H2B.8 forms chromatin condensates by increasing interactions between H2B.8-enriched chromosomal regions (Figs. 2C, 3C and 6, B and C). Due to H2B.8 deposition within AT-rich sequences in both pericentromeric regions and chromosomal arms (Fig. 4E and figS. S4A), broad chromosomal regions are concentrated by phase separation. In interphase somatic cells, euchromatin takes up most of the nuclear volume (*52*). Because H2B.8 is abundant in euchromatin, H2B.8-induced chromatin condensation is highly effective at condensing nuclei (Figs. 2, A and D).

Despite its effectiveness, H2B.8 compacts nuclei without compromising transcription (Figs. 5, A and B). For species with swimming sperm, in which the size of the sperm head is crucial (*1, 2, 10*), DNA condensation may be paramount and transcription dispensable. Protamines can greatly condense DNA and have been identified in the sperm of almost all multicellular eukaryotic lineages except angiosperms (flowering plants) and gymnosperms (Fig. 3A). This includes bryophytes and pteridophytes, both of which have motile sperm and use protamines for sperm condensation (Fig. 3A) (*11–15, 24*). In contrast, H2B.8 is specific to flowering plants. Flowering plants have immotile sperm and may benefit from a less radical approach that condenses nuclei without limiting transcription. We speculate that fertilization, which takes place at ovules deeply embedded in angiosperm maternal tissues, might favor smaller sperm nuclei. Consistent with this idea, gymnosperms, which have exposed ovules, produce sperm with uncondensed nuclei (*12*) and lack H2B.8 (Fig. 3A). Therefore, H2B.8 is an angiosperm evolutionary innovation that achieves a moderate level of chromatin condensation compatible with active transcription.

## ACKNOWLEDGMENTS

We thank C. Dean, S. Penfield and D. Zilberman for constructive comments on the manuscript. We also thank the John Innes Centre Bioimaging Facility (S. Lopez and E. Wegel) for assistance with microscopy, the Norwich BioScience Institute Partnership Computing infrastructure for Science Group for High Performance Computing resources, and Annoroad Gene Technology Co., Ltd (Beijing, China) for Hi-C library preparation and sequencing.

## Author contributions

TB, SH and XF conceived the study, designed the experiments and interpreted the results. TB, SH, LW, SZ, LS, and GS performed experiments. TB, SH, SZ, MV, and XF performed data analysis. TB, SH and XF wrote the manuscript, and all authors commented on the manuscript.

## Data and materials availability

Sequencing data generated in this study (ChIP-seq, RNA-seq and Hi-C) have been deposited in the Gene Expression Omnibus (GEO) under accession no. GSE161366. All remaining data are in the main paper or the supplementary materials. Further information and requests for resources and reagents should be directed to Xiaoqi Feng.

## Materials and Methods

### Plant growth conditions

*Arabidopsis thaliana* plants (Col-0 ecotype) used in this study were grown under long day (16 hr light, 8 hr dark) conditions at 22 °C and 70% humidity. Seedlings were grown on germination medium (GM) plates without glucose under the same conditions.

### Generation of CRISPR-Cas9 mutants

Mutant alleles of *H2B.8* were generated by CRISPR-Cas9 (*53*). Four sgRNAs (Table S2) were designed using CHOPCHOP-v3 (*54*) and cloned using the Golden Gate system (*53*). Constructs were transformed via *Agrobacterium tumefaciens* strain GV3101 using floral dip to WT Col-0 *Arabidopsis thaliana* (*55*). Transformants were screened by Sanger sequencing. Selected lines were taken to the next generation to produce homozygous mutants without the Cas9. Line *h2b.8-1* was genotyped by dCAPS with EcoNI, whilst line *h2b.8-2* was genotyped by PCR (Table S2).

### Reporter and ectopic expression construct cloning

For the H2B.8 reporter constructs (*pH2B.8::H2B.8-eGFP* and *pH2B.8::H2B.8-Myc*), approximately 2 kb upstream of H2B.8 was cloned as the promoter and the H2B.8 gDNA sequence was amplified (Table S2). Using MultiSite Gateway Technology (Thermo Fisher Scientific), the PCR products were ligated to P4P1r and pDONR207, respectively. Sequences were assembled to the expression vector pK7m34GW with a C-terminal eGFP or 3xMyc tag in P2rP3. The ectopic H2B.8 expression vector (*p35S::H2B.8-eGFP*) was generated in the same way, using the 35S promoter.

Ectopic H2B.8ΔIDR expression construct (*p35S::H2B.8ΔIDR-YFP*) was generated by overlapping PCR to remove the IDR sequence whilst ectopic H2B.2 (*p35S::H2B.2-YFP*) was cloned from gDNA (Table S2). Products were ligated to pCAMBIA1300 vector backbone containing the 35S promoter and a C-terminal YFP using the In-Fusion cloning system (Takara Bio).

The H2B.8 reporter constructs (*pH2B.8::H2B.8-eGFP* and *pH2B.8::H2B.8-Myc*) were transformed to *h2b.8-1* mutant plants of T3 generation. Ectopic expression constructs (*p35S::H2B.8-eGFP*, *p35S::H2B.8ΔIDR-YFP* and *p35S::H2B.2-YFP*) were transformed to the wild type (WT). Single insertion transgenic lines in T3 or T4 generations were used in this study.

### Sperm and vegetative nuclei total protein extraction

Sperm and vegetative nuclei were isolated by FACS (Fluorescence-Activated Cell Sorting), as previously described (*56*). Nuclei were pooled and 0.45 volumes of 3.2 x lysis buffer (10% SDS, 100 mM TEAB, pH 7.55) was added. Nuclei were lysed at 95 °C for 5 min, then centrifuged at 13000 x g for 8 min at RT. Lysate was moved to a new tube. One tenth volume 12% phosphoric acid was added and mixed by pipetting. Then, six times volumes of S-Trap buffer (90% aqueous MeOH, 100 mM TEAB, pH 7.1) was added and mixed by pipetting. Protein was loaded to an S-Trap Micro column (Protifi) by centrifugation, 4000 x g for 30 s. The column was washed three times with 150 µl S-Trap buffer. Protein was digested on column with 4 µg trypsin in 50 mM TEAB at 47 °C for 1 hr. Peptides were eluted sequentially by centrifugation (4000 x g for 30 s) with 40 µl 50 mM TEAB, 40 µl 0.2% formic acid and 35 µl 50% ACN 0.2% formic acid.

### Liquid Chromatography–Mass Spectrometry (LC-MS)

The eluted peptide solutions were dried down, and the peptides dissolved in 0.1% TFA / 3% acetonitrile for liquid chromatography with tandem mass spectrometry. Sperm and vegetative nuclei samples were analyzed by nanoLC-MS/MS on an Orbitrap Fusion Tribrid mass spectrometer coupled to an UltiMate 3000 RSLCnano LC system (Thermo Fisher Scientific). The samples were loaded and trapped using a pre-column with 0.1% TFA at 20 µl/min for 3 min. The trap column was then switched in-line with the analytical column (nanoEase M/Z column, HSS C18 T3, 100 Å, 1.8 µm, Waters) for separation using the following long gradient of solvents A (water, 0.05% formic acid) and B (80% acetonitrile, 0.05% formic acid) at a flow rate of 0.3 µl/min: 0-3 min 3% B (trap only); 3-14 min linear increase B to 13%; 14-113 min increase B to 39%; 113-123 min increase B to 55%; followed by a ramp to 99% B and re-equilibration to 3% B. Data were acquired with the following mass spectrometer settings in positive ion mode: MS1/OT: resolution 120K, profile mode, mass range *m/z* 300-1800, AGC 4e^5^, fill time 50 ms; MS2/IT: data dependent analysis was performed using HCD fragmentation with the following parameters: top30 in IT rapid, centroid mode, isolation window 1.6 Da, charge states 2-5, threshold 1.9e^4^, CE = 30, AGC target 1.9e^4^, max. inject time 35 ms, dynamic exclusion 1 count, 15 s exclusion, exclusion mass window ±5 ppm.

For sperm and vegetative nuclear proteomes, recalibrated peak lists were generated with MaxQuant-1.6.1.0 (*57*) in LFQ mode using the TAIR10_pep_20101214 *Arabidopsis* protein sequence database (TAIR, 35386 entries) plus the MaxQuant contaminants database (245 entries). The quantitative LFQ results from MaxQuant with default parameters were used together with search results from an in-house Mascot Server 2.4.1 (Matrix Science) on the same databases. For all searches, a precursor tolerance of 6 ppm and a fragment tolerance of 0.6 Da was used. The enzyme was set to trypsin/P with a maximum of 2 allowed missed cleavages; oxidation (M) and acetylation (protein N-term) were set as variable modifications; carbamido-methylation (C) as fixed modification. The search results were imported into Scaffold 4 (Proteome Software) using identification probabilities of 99% for proteins and 95% for peptides.

### Histone alignments and disorder predictions

Alignments of histone DNA and protein sequences was performed using CLC Main Workbench software version 8.1 (QIAGEN). Predictions of intrinsic disorder were undertaken by PONDR (*58*), using the VL-XT algorithm. Raw data was plotted using ggplot2 in R-3.6.0 (*59, 60*).

### Histone H2B phylogenetic analysis

Plant H2B protein sequences were downloaded from Phytozome (*61*), Congenie (*62*), Waterlily Pond (*63*), Magnoliid genomes (*64, 65*) and Uniprot (*66*). Human and yeast H2B sequences were obtained from Uniprot and used as out-groups for phylogenetics.

Sequences were imported to MEGA-X (*67*) and aligned using MUSCLE with default parameters. The phylogeny was generated using Neighbor-Joining testing, applying the Poisson model, and allowing for uniform substitution rates. H2B.8 homologs were identified owing to the distinct branch formed, separate from canonical H2B variants. Several representative H2B.8 homologs were searched using BLAST (*68*) to ask whether such homologs are specific to flowering plants. H2B.8 homologs are presented in Table S1.

### Confocal microscopy and analysis

Microspores and pollen were isolated as described previously (*56*), stained with Hoechst 33342, and examined under a Leica SP8X confocal microscope. Young embryos were dissected (*69*) and stained with propidium iodide for imaging. Mature embryos were isolated from dry seeds using a stereo microscope. Mature embryos and seedlings (including roots) were stained in PBS with 0.1% Triton X-100 and 0.5 µg/ml DAPI for 5-10 min before microscopic examination (Zeiss 880, Airyscan mode). Immunofluorescence was performed with two-week-old seedlings as described previously (*23*).

Sperm nuclear size was quantified from DAPI-stained whole pollen confocal images using a semi-automated pipeline in ImageJ adapted from (*70*). Briefly, auto-threshold was used to obtain nuclei and then processed using Guassian blur to smooth edges. The auto-threshold was repeated and then nuclei were selected using the wand tool. Measurements were then obtained for nuclear area (μm^2^). Somatic nuclei selected for analysis were vascular cylinder cells in the elongation zone of the root tip. Such nuclei were selected owing to the ability to accurately identify the cell type within the tissue. Using ImageJ, Z-stacks were divided into substacks of different cell layers within the root tip. Maximum intensity projections were then obtained to account for slight differences in the depth of nuclei. Images were analyzed in the same semi-automated way as per sperm nuclei. Statistical analysis was undertaken in R; we used ANOVA followed by Tukey’s post-hoc test for pairwise comparisons.

Classification of H3K9me2 foci was undertaken as previously described (*23*). H2B.8-mediated chromatin aggregates were classified in the same way.

To quantify size and number of H2B.8 aggregates and H3K9me2 domains, we used the 3D ImageJ Suite tools (*71*). In the according fluorescence channel, we undertook 3D Nuclei Segmentation using Otsu thresholding (*71*). The number and volumes of segments were plotted in R using ggplot2.

### 3D Structured Illumination Microscopy

Sperm and vegetative nuclei were isolated from pollen as described previously (*56*) and resuspended in 200 μl Galbraith buffer (45 mM MgCl_2_, 30 mM sodium citrate, 20 mm MOPS, 0.1% Triton X-100, pH 7.0). Nuclei were extracted from seedlings by finely chopping with a razor blade in lysis buffer (15 mM Tris pH 7.5, 2 mM EDTA, 0.5 mM spermine, 80 mM KCl, 20 mM NaCl, 0.1% Triton X-100). Suspension was filtered through a 35 μm filter (Corning) to a 1.7 ml tube. Nuclei were pelleted at 500 x g for 3 mins and resuspended in 200 μl lysis buffer.

Nuclei were fixed in solution with 4% MeOH-free formaldehyde (Thermo Scientific) for 5 mins. HiQA No. 1.5H coverslips (CellPath) were washed with 10% HCl for 30 mins and then washed three times in H_2_O for 5 mins to remove impurities. Fixed nuclei were spun onto coverslips at 500 x g for 3 mins using a Shandon Cytospin 2. Fixation was repeated by blotting nuclei with 4% MeOH-free formaldehyde for 5 mins. Fixative was removed and coverslips were washed three times in PBS, 5 mins per wash. Nuclei were blocked with 3% BSA in PBS with 0.1% Tween-20 (PBST) for 30 min in a humidified chamber. If performing immunostaining, antibodies were diluted 200-fold in 3% BSA in PBST and then blotted to nuclei on coverslips. Antibody incubation occurred overnight at 4 °C. Primary antibodies were removed by washing three times with PBST for 5 mins. Secondary antibodies were diluted similarly to primary and then added to nuclei. Incubation occurred for 1 hr at RT in a humidified chamber. If not performing immunostaining, the protocol resumes at this point. PBST washes were repeated as before. Nuclei were stained in the dark with either DAPI or SYBR Green (Invitrogen) at 2 mg/μl or 100 x dilution, respectively for 5 mins. DNA stain was removed by washing in H_2_O for 5 mins. Coverslips were adhered to slides in 13 μl VECTASHIELD H-1000 mounting medium. Nuclei were imaged using a 63x oil immersion lens on a Zeiss Elyra PS.1 super-resolution microscope.

Three dimensional reconstructions for SIM were undertaken using Zeiss Zen Black software. Intensity profiles associated with images were acquired using the Interactive 3D Surface Plot plugin for ImageJ (*72, 73*).

To acquire voxel intensities from WT and *h2b.8* sperm, nuclei were segmented using Otsu thresholding and individual voxels were extracted. Fluorescence intensity was normalized by dividing by total nuclear intensity. Density of binned voxel intensities were plotted with ggplot2 in R.

### Histone purification from *E. coli*

Sequences for H2B.8 and H2B.2 were cloned into the pET28a+ vector with a non-cleavable C-terminal 8×His-tag and then transformed to *Escherichia coli* strain BL21 (Tiangen). Cells were grown to OD 0.8 at 37 °C in LB media with 30 μg/ml kanamycin. Histone expression was induced by addition of 0.1 mM isopropyl-β-d-thiogalactopyranoside (IPTG) and incubation overnight at 16 ℃. Cells were collected by centrifugation at 4000 rpm for 20 min and resuspended in lysis buffer (20 mM Tris-HCl, 500 mM NaCl, pH 8.0). Cells were lysed by ultrasonication and debris was pelleted at 20000 x g. The supernatant was applied to a 5 ml HisTrap HP column (Cytiva) on AKTA pure (Cytiva). Target proteins were eluted at ∼250 mM imidazole concentration during gradient elution. The peaks eluted were applied to a Superdex 200 Increase 3.2/300 (Cytiva) gel filtration column, then dialyzed and concentrated using in vitro phase separation assay buffer (20 mM HEPES, 150 mM NaCl, pH 8.0).

The H2B.8ΔIDR sequence was cloned to a modified pET11 expression vector (Novagen) as previously described (*74*). The expression vector contains a solubility MBP tag followed by a TEV cleavage site and a GFP tag upstream of the insertion site and a non-cleavable C-terminal 8×His-tag at downstream of the insertion site. Proteins were expressed and lysed as before, besides using 50 μg/ml ampicillin for selection. Purification was performed as previously, with target proteins between 300 mM and 500 mM imidazole concentration during gradient elution. For *in vitro* phase separation assays, the MBP tag was cleaved by incubating with ∼0.02 mg/ml 6×Histag-TEV protease overnight at 4 °C, and the cleaved GFP-H2B.8ΔIDR was tested by western blot using His-tag antibody (Huaxingbio).

### *In vitro* phase separations assays

All *in vitro* experiments were performed in phase separation assay buffer (20 mM HEPES, 100 mM NaCl, pH 7.4). *In vitro* experiments were recorded on 384 low-binding multi-well 0.17 mm microscopy plates (In Vitro Scientific) and sealed with optically clear adhesive film. Imaging was performed with a NIKON A1 microscope equipped with a 100x oil immersion objective. NIS-Elements AR Analysis was used to examine images.

### *In vitro* and *in vivo* FRAP

*In vitro* FRAP (Fluorescence Recovery After Photobleaching) experiments were carried out with a NIKON A1 microscope equipped with a 100x oil immersion objective. Droplets were bleached with a 488- or 561-nm laser pulse (3 repeats, 70% intensity, dwell time 1 s).

Root nuclei expressing H2B.8 (*p35S::H2B.8-eGFP*) were imaged using Airyscan mode with a Zeiss 880 confocal microscope. Individual H2B.8 foci were photobleached and imaged at intervals over time.

Images were processed using the ImageJ StackReg plugin (*72*). Post-bleach intensity was normalized to pre-bleach levels to obtain a measure of recovery. Data was plotted using ggplot2 in R.

### Pollen and seedling native ChIP-seq library preparation, sequencing and analysis

Ten-day-old seedlings were ground with a pestle and mortar in liquid N_2_ and homogenized in nuclei isolation buffer (0.25 M sucrose, 15 mM PIPES pH 6.8, 5 mM MgCl_2_, 60 mM KCl, 15 mM NaCl, 1 mM CaCl_2_, 0.9% Triton X-100, 1 mM PMSF, 1x Protease Inhibitor Cocktail (Roche)) for 15 min. Nuclei were separated from debris by filtering through two layers of miracloth (Merck Millipore). For pollen nuclei, we collected open flowers and isolated pollen in Galbraith buffer. Nuclei were released by vertexing pollen with glass beads twice in nuclei isolation buffer. The homogenate was filtered through 40 µm and 10 µm cell strainers successively to obtain nuclei.

Nuclei suspension from seedlings or pollen was centrifuged at 4000 x g for 10 min and pellets were resuspended in TM2 (50 mM Tris-HCl, 2 mM MgCl_2_, 0.25 M sucrose, 1 mM PMSF, 1 x Protease Inhibitor Cocktail). After cold centrifugation at 4000 x g for 5 min, nuclei were resuspended in MNase digestion buffer (50mM Tris-HCl pH 7.5, 5mM CaCl_2_, 0.25 M sucrose, 1 mM PMSF, 1 x Protease Inhibitor Cocktail) with appropriate amount of MNase (New England Biolabs) and incubated at 37 °C for 10 min. Digestion was stopped by adding EDTA to a final concentration of 25 mM. One tenth volume of 1% Triton X-100 and 1% sodium deoxycholate was added and left on ice for 15 min. Then, the reaction was diluted by adding low salt buffer (50 mM Tris-HCL pH 7.5, 10 mM EDTA, 150 mM NaCl, 0.1% Triton X-100, 1 mM PMSF, 1 x Protease Inhibitor Cocktail) and rotated for 1 hr at 4 °C. After centrifugation, the supernatant was used for immunoprecipitation with pre-washed GFP-Trap beads (Chromotek) overnight at 4 °C. Beads were washed twice each with low salt buffer and high salt buffer (50 mM Tris-HCL pH 7.5, 10 mM EDTA, 300 mM NaCl, 0.1% Triton X-100, 1 mM PMSF) and eluted in elution buffer (0.1 M NaHCO_3_, 1% SDS) by shaking at 65 °C for 15 min. The eluates were digested with Proteinase K and RNase A before phenol-chloroform DNA extraction. Libraries were prepared using Ovation Ultralow System V2 and sequenced on NextSeq 500 (Illumina) with 2 × 38 bp paired end reads.

Sequencing reads were mapped to TAIR10 with Bowtie2-2.3.4.1 (*75*), retaining mononucleosomal fragments. Bigwig files were generated by normalizing IP bam files to respective inputs using deepTools-3.1.1 (*76*). Two replicates for each experiment were confirmed to be highly correlated; a single replicate was used for downstream analyses. Profiles were visualized using IGV-2.6.2 (*77*). Data underlying metaplots and heatmaps were generated with deepTools and plotted with a custom script in R.

TE classes were defined by seedling H3K9me2 enrichment (log_2_(IP/input)). Considering the bimodal distribution of H3K9me2 enrichment at TEs, 0.7 was selected as the cut-off for heterochromatic (> 0.7) and euchromatic (< 0.7) classes.

To generate peaks, H2B.8 enrichment was calculated over 50 bp windows and those with > 1.2 log_2_(IP/input) were retained. Windows within 150 bp were merged using BEDtools-2.28.0 (*78*). Regions were filtered by size, with those < 200 bp removed from analysis. H2B.8 enrichment was then calculated over the new regions, those with < 1.2 log_2_(IP/input) were discarded. The remaining regions were defined as H2B.8 peaks.

For genome coverage, peaks were divided into 50 bp windows and partitioned into gene, TE or intergenic groups depending on overlaps. Overlaps with genes and TEs for volcano plots were determined using BEDtools; 25% of the feature was required to be covered by a peak to be defined as an overlap.

Downloaded data (*79–86*) was mapped and prepared in the same way. Principal Component Analysis and Spearman’s rank correlations were calculated with deepTools and plotted in R with ggplot2.

### Bisulfite-seq analysis

Downloaded sequencing reads (*87*) were processed using TrimGalore-0.4.1 (https://github.com/FelixKrueger/TrimGalore) with default parameters. Reads were mapped to TAIR10 using Bismark-0.22.2 (*88*) and methylation was called using MethylDackel-0.5.2 (https://github.com/dpryan79/MethylDackel), selecting --CHG and --CHH options. CG methylation data was used in PCA.

### Sperm cell and seedling RNA-seq library preparation, sequencing and analysis

Sperm cells were isolated by FACS as described previously (*89*). RNA was extracted with Direct-zol RNA Microprep. Plant RNeasy Mini Kit was used to extract RNA from 10-day-old seedlings. Libraries were prepared using a Universal RNA-Seq library preparation kit and sequenced on NextSeq 500 (Illumina) with single end (76 bp) or paired end (2 × 38 bp) reads.

Sequencing reads were mapped to TAIR10 with TopHat-2.0.10 (*90*). Kallisto-0.43.0 (*91*) and Sleuth-0.30.0 (*92*) were used to obtain TPM and *q*-values, respectively. Differentially expressed genes and TEs were identified by | log_2_ (TPM fold change) | ≥ 2 or ≤ -2 and *q* < 0.05. Volcano plots were generated with a custom ggplot2 R script.

Downloaded data (*93–95*) was mapped and TPM values were obtained in the same way.

### Hi-C library preparation, sequencing and analysis

10-day-old seedlings were harvested and fixed with 20 ml 2% formaldehyde solution for 15 min in vacuum conditions at room temperature and then quenched by adding 2.162 ml 2.5 M glycine. Fixed seedling tissue was rinsed with water three times and dried with tissue paper.

The nuclei were released by grinding in liquid nitrogen and then resuspended with 25 ml of extraction buffer I (0.4 M sucrose, 10 mM Tris HCl pH 8, 10 mM MgCl_2_, 5 mM β-mercaptoethanol, 0.1 mM PMSF, 13 μl protease inhibitor). Nuclei were filtered through miracloth (Calbiochem) and then centrifuged at 4000 rpm for 20 min at 4 °C. The supernatant was discarded whilst the pellet was resuspended with 1 ml of extraction buffer II (0.25 M sucrose, 10 mM Tris HCl pH 8, 10 mM MgCl_2_, 1% Triton X-100, 5mM β-mercaptoethanol, 0.1mM PMSF, 13 μl protease inhibitor). Then, the mixture was centrifuged at 14000 rpm for 10 min at 4 °C and the pellet was resuspended with 300 μl of extraction buffer III (1.7 M sucrose, 10 mM Tris HCl pH 8, 0.15% Triton X-100, 2 mM MgCl_2_, 5 mM β-mercaptoethanol, 0.1 mM PMSF, 1 μl protease inhibitor). The mixture was then loaded onto an equal amount of clean extraction buffer III and centrifuge at 14000 rpm for 10 min. The pelleted nuclei were washed twice with 1x ice cold CutSmart buffer and finally resuspended in 0.5 ml volume. SDS was applied to permeabilize nuclei at 65 °C for 10 min, Triton X-100 was added to quench SDS. Thereafter, chromatin was digested with 400 units MboI overnight at 37 °C with gentle rocking. MboI was then denatured to cease activity.

Digested chromatin underwent DNA end repair with biotin-14-dCTP insertion followed by blunt-end ligation. After decrosslinking with proteinase K at 65 °C, DNA was purified by phenol chloroform extraction method. Biotin-14-dCTP was removed from non-ligated DNA fragment ends using T4 DNA polymerase. DNA was sheared to a range of 200 to 600 bp by sonication. Next, the fragments underwent end repair and were pulled down by streptavidin C1 magnetic beads to enrich for fragments containing contact information. Fragment ends were then A-tailed, sequencing adapters were ligated, and libraries were amplified by PCR for 12-14 cycles. Following purification, libraries were sequenced using the Illumina HiSeq X Ten platform with 2×150 bp length reads. The Hi-C library construction and sequencing were conducted by Annoroad Gene Technology Co., Ltd (Beijing, China).

Sequencing reads were mapped to the TAIR10 reference genome using the HiC-Pro-2.11.1 pipeline (*96*). The bam files (bwt2merged.bam) generated by HiC-Pro containing with mapped reads were used as input files for FAN-C-0.9.8 (*97*). The module ‘fanc auto’ was applied to generate 500 kb, 100 kb, 50 kb, 10 kb, 1kb contact matrices (hic files). The resultant hic files with 100 kb resolution were directed to the ‘fanc expected’ module to calculate the expected interaction probability against genomic distance for intrachromosomal interaction. For matrix and score comparison, the default comparison method of fold-change was employed by ‘fanc compare’ command. The outputs (hic object) were transferred to text files by ‘fanc dump’ and were visualized as heatmaps in R using ggplot2. To explore whether higher contacts observed in *p35S::H2B.8-eGFP* depends on H2B.8 incorporation, the genome was binned into 1 kb windows. H2B.8 signals (log_2_(IP/input)) of each window were generated and sorted into 20 quantiles by strength. The interaction frequency differences (values generated by FAN-C at 1 kb resolution) of each quantile pair for either short-range interactions or interactions between pericentromeric regions and chromosome arms were averaged and plotted as heatmaps. The resolution of our Hi-C data was estimated as previously reported (*98*). Our Hi-C data was deemed to achieve 1 kb resolution as 80% of genomic bins (1 kb) had >1000 contacts. Our WT data was compared to published contact matrices (*47, 48*).

### Plots and statistics

Statistical tests performed on experimental data and sample sizes are noted in figure legends. Boxplots show median (thick black bar) and first and third quartiles, with lower and upper whiskers extending to 1.5 times the interquartile range of the first and third quartiles or the highest and lowest values, respectively.

---

**Fig. S1.**
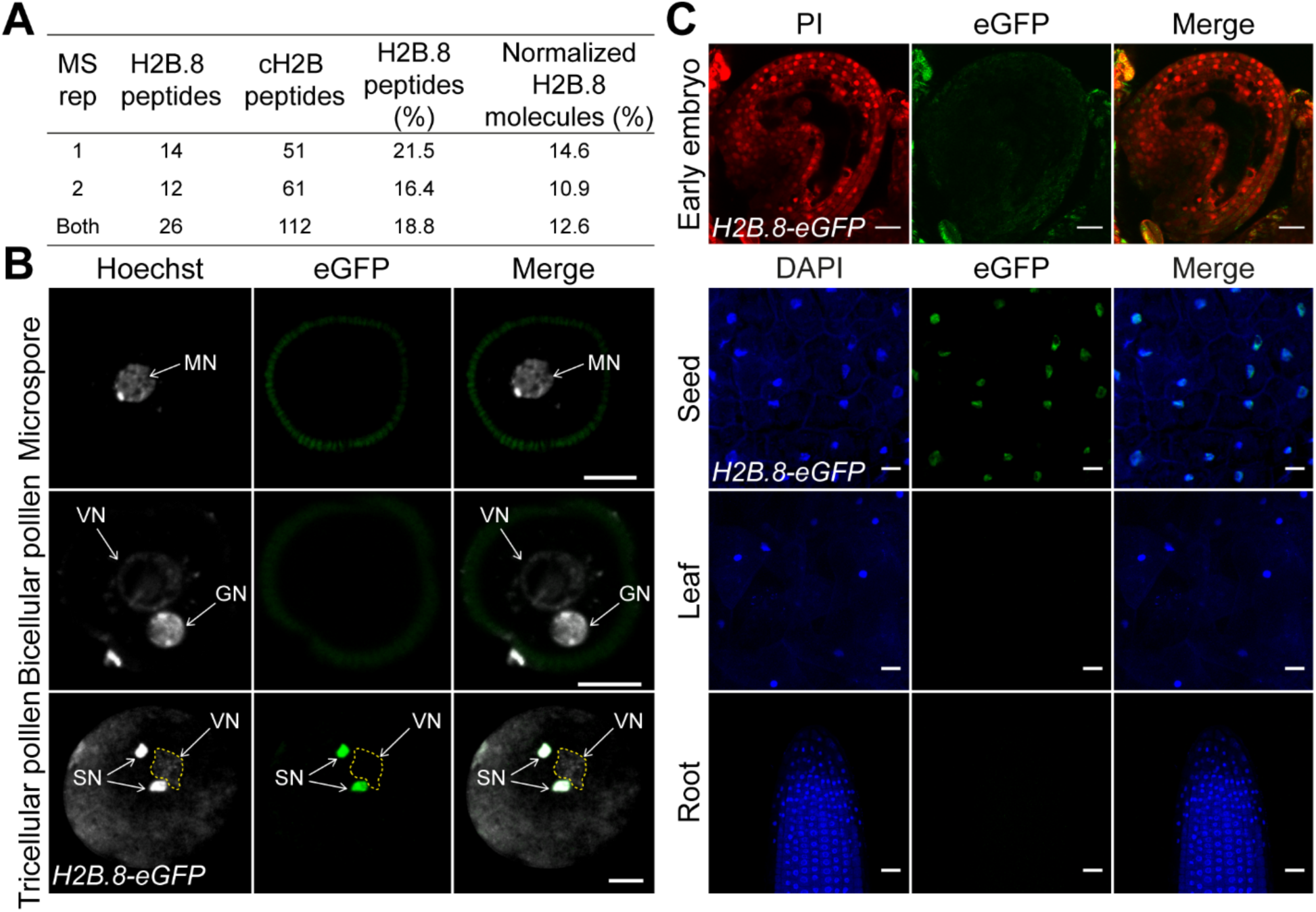
H2B.8 is specifically expressed in sperm and mature seeds. (**A**) Peptide counts from two biological replicates of *Arabidopsis* sperm nuclei mass spectrometry. H2B.8 is compared to canonical H2B (cH2B) peptide counts. Normalized proportion of total H2B accounts for larger size of H2B.8 compared to canonical variants (243 amino acids versus ∼151). (**B**) Confocal images of H2B.8 (*pH2B.8::H2B.8-eGFP*) incorporation through male gametogenesis. Lowest panel is a duplicate of Fig. 1D. MN, VN, GN and SN, respectively, microspore, vegetative, generative and sperm nucleus. Scale bars, 5 μm. (**C**) Confocal images of H2B.8 (*pH2B.8::H2B.8-eGFP*) in indicated tissues. Scale bars, 20 μm (2-cell embryo, leaf, and root) and 5 μm (seed).

**Fig. S2.**
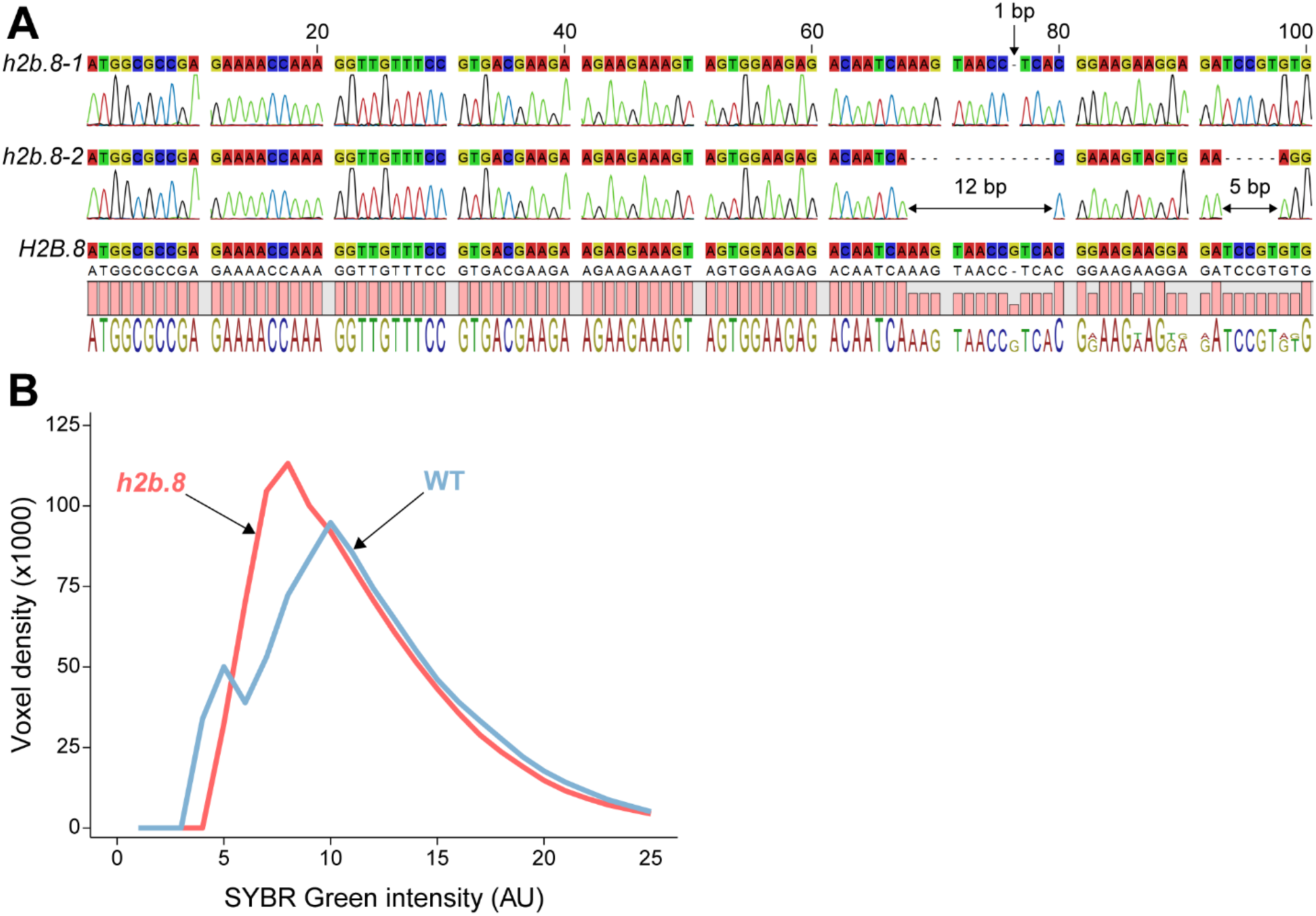
H2B.8 is required for sperm chromatin aggregation. (**A**) Alignment of *h2b.8* CRISPR lines. *h2b.8-1* has a single base deletion at 76 bp as indicated by an arrow, leading to a premature stop codon after 33 amino acids. *h2b.8-2* has a 12 bp deletion after 67 bp and another of 5 bp after 92 bp, producing a truncated 54 amino acid protein. (**B**) Density plot of individual SYBR Green-stained voxel intensities from wild-type (WT, blue) and *h2b.8* (red) sperm nuclei. n = 30 nuclei for WT and *h2b.8*.

**Fig. S3.**
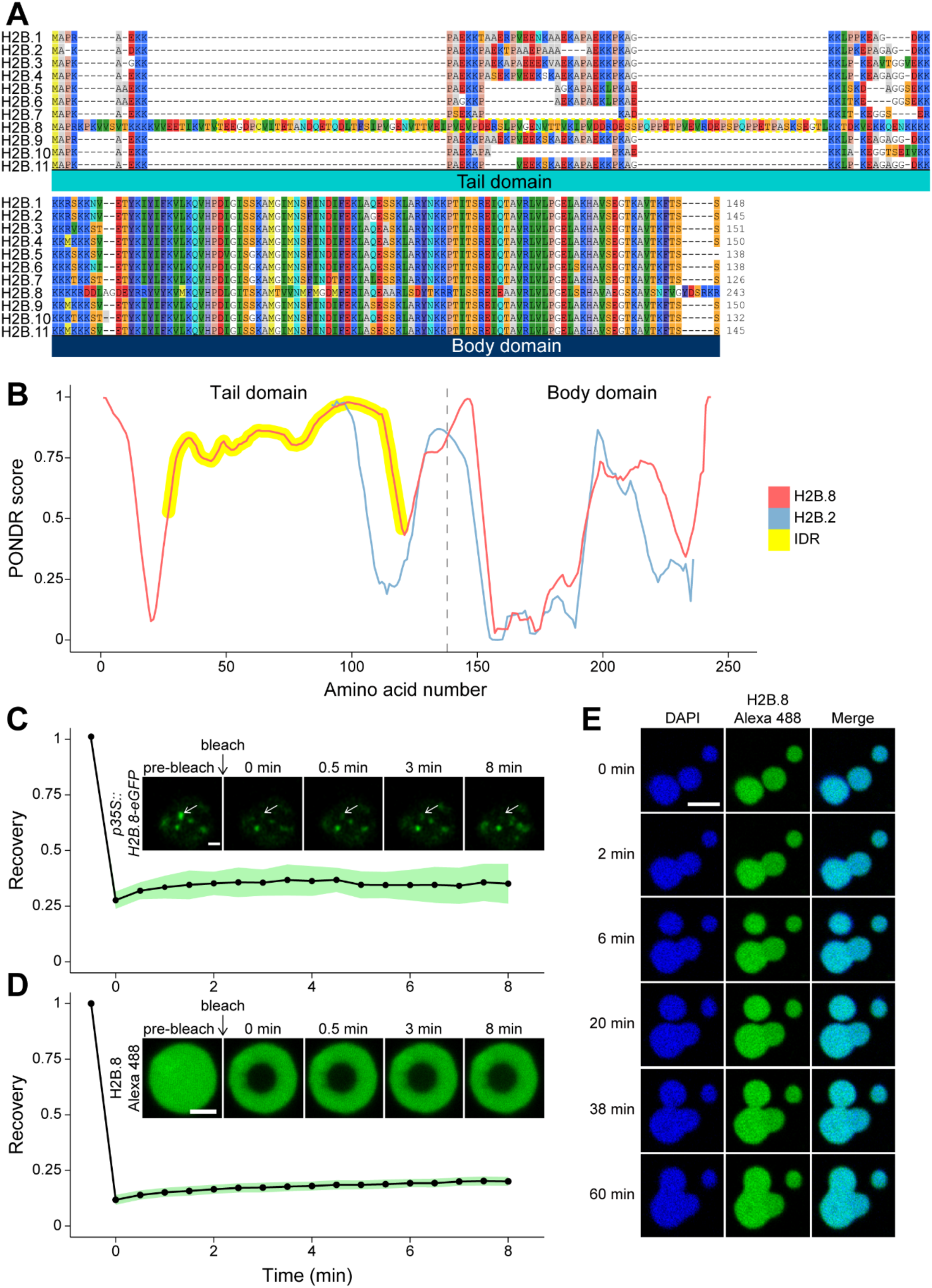
Phase separation property of H2B.8. (**A**) Alignment of *Arabidopsis* H2B variants (N-terminal tail – upper panel, C-terminal body – lower panel). The N-terminal tail IDR of H2B.8 is highlighted with a yellow dashed box. Amino acids are colored according to the RasMol scheme. (**B**) Intrinsic disorder prediction by PONDR of H2B.8 (red) and H2B.2, a canonical H2B (blue). Histone profiles are aligned at the interchange between tail and body domains (dashed line). H2B.8 N-terminal tail IDR is highlighted in yellow. (**C**) FRAP trace of *in vivo* H2B.8 (*p35S::H2B.8-eGFP*) chromatin aggregates. n = 11 foci. Green area indicates standard deviation. Scale bar, 2 μm. (**D**) FRAP trace of *in vitro* H2B.8 condensate. n = 8 foci. Green area indicates standard deviation. Scale bar, 2 μm. (**E**) *In vitro* H2B.8 condensate fusion over time. Scale bar, 5 μm.

**Fig. S4.**
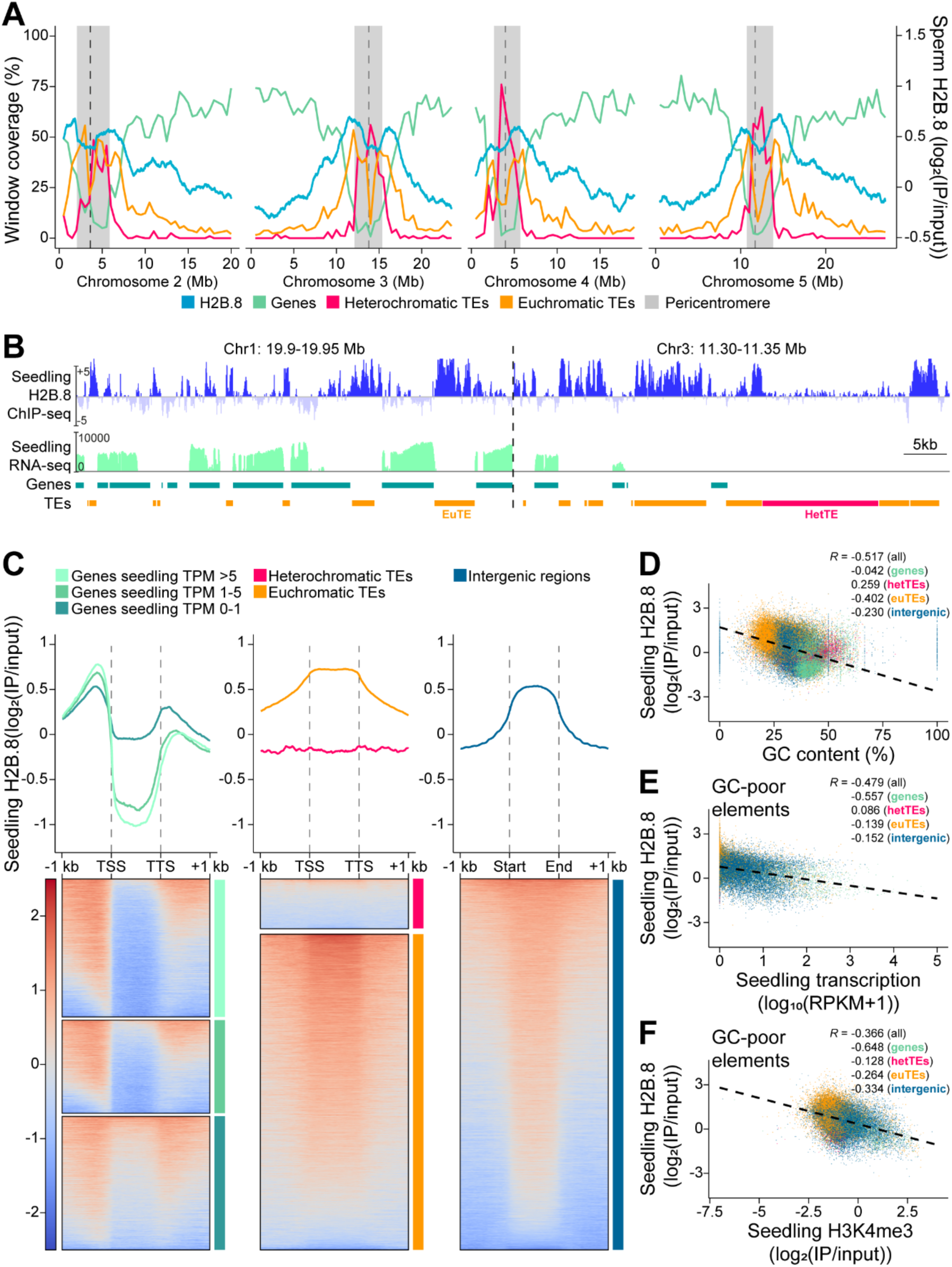
Ectopically expressed H2B.8 in seedlings exhibits a similar deposition profile as native H2B.8 in sperm. (**A**) Coverage of genes, euchromatic TEs and heterochromatic TEs (left Y axis; 500 kb windows) and H2B.8 enrichment in sperm (right Y axis; 1 kb windows) along Chromosomes 2 to 5. Chromosome 1 is shown in Figure 4E. (**B**) Genome snapshots of H2B.8 abundance in seedlings (*p35S::H2B.8-eGFP*; log_2_(IP/input)), seedling transcription (log_2_(RPKM), and gene and TE annotations (orange, euchromatic TE; magenta, heterochromatic TE) over representative 50 kb regions. RPKM, Reads Per Kilobase of transcript per Million mapped reads. (**C**) Profiles and associated heatmaps of seedling *p35S::H2B.8-eGFP* enrichment over genes (grouped by seedling expression), TEs (grouped by chromatin state) and intergenic regions. (**D**) Scatterplot showing anticorrelation of ectopic H2B.8 enrichment in seedling with GC content (%) over denoted genomic features. *R*, Spearman’s Rank. (**E** and **F**) Scatterplots illustrating anticorrelation of ectopic H2B.8 enrichment in seedling with seedling transcription (E) or H3K4me3 (F) over genomic features with low (<34.5%) GC content. *R*, Spearman’s Rank.

**Fig. S5.**
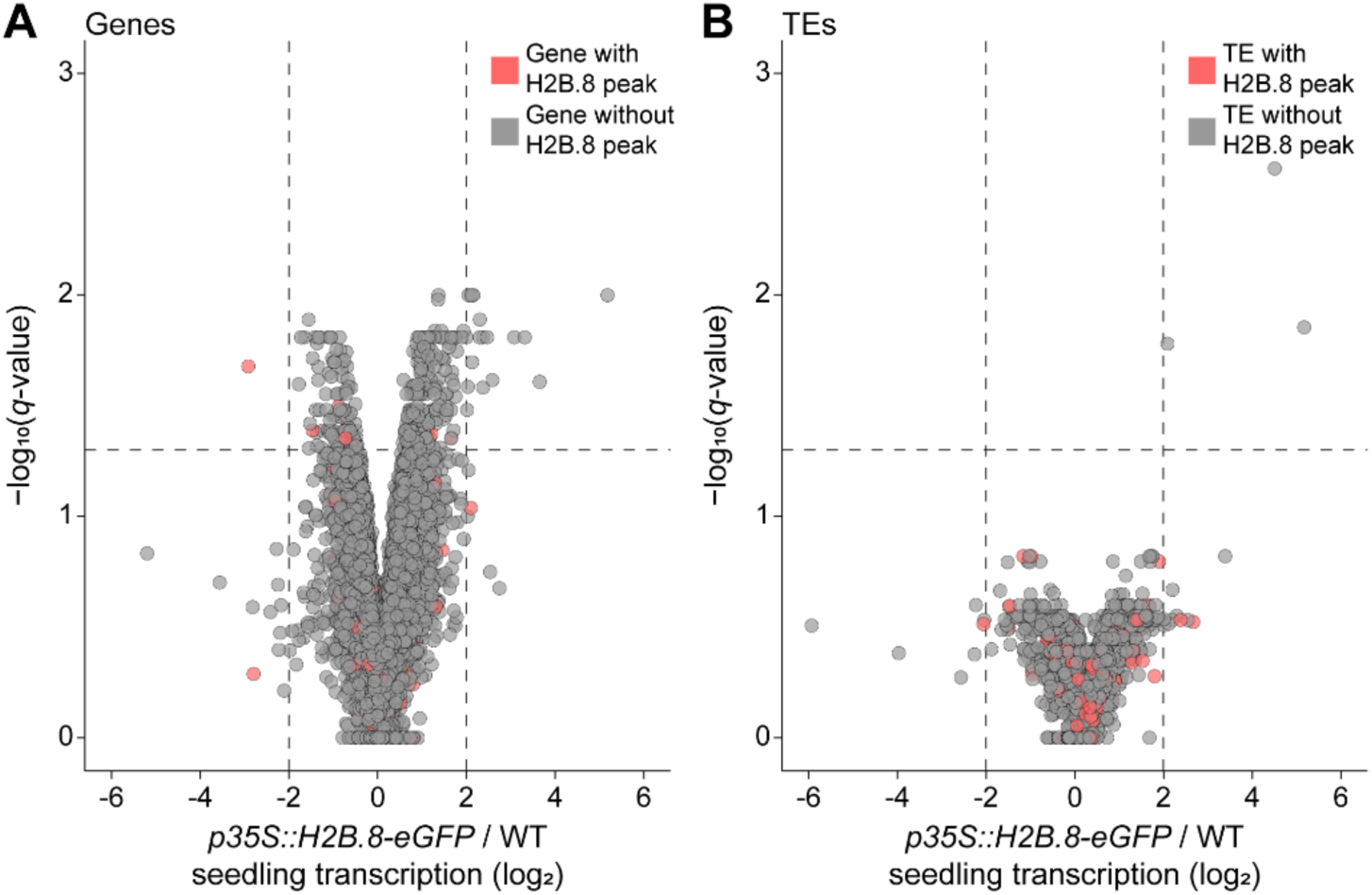
Ectopically expressed H2B.8 has a negligible effect on seedling transcription. (**A** and **B**) Volcano plots showing differential gene (A) or TE (B) expression between *p35S::H2B.8-eGFP* and wild-type (WT) seedlings. Differentially expressed genes/TEs were defined as log_2_(*p35S::H2B.8-eGFP* versus WT TPM fold change) ≥2 or ≤-2 and *q* < 0.05 (likelihood-ratio test). TPM, Transcripts Per Million. n = 20285 (A) and 1472 (B).

**Fig. S6.**
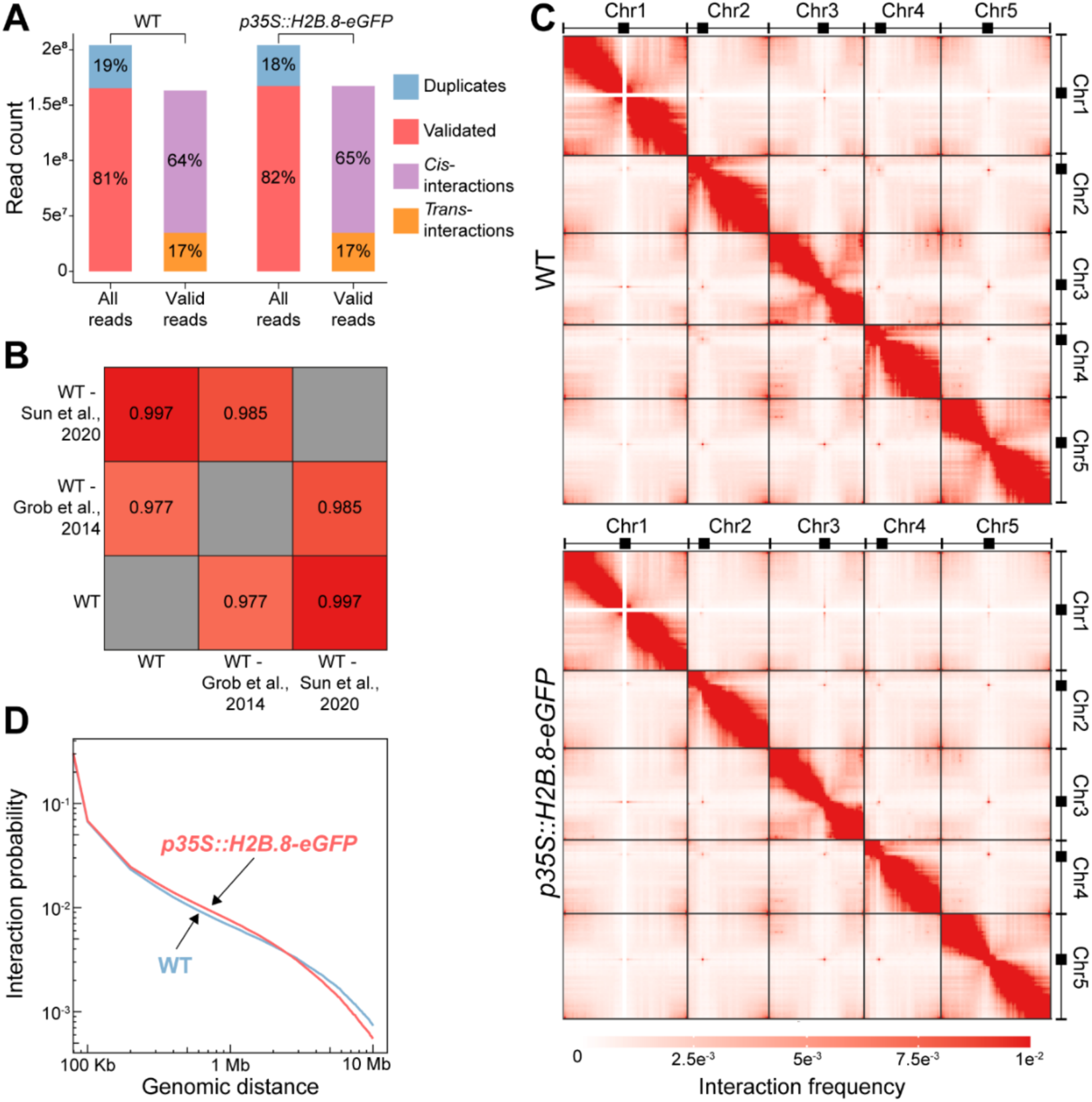
H2B.8 affects intra- and inter-chromosomal interactions. (**A**) Sequencing read distributions of wild-type (WT) and *p35S::H2B.8-eGFP* seedling Hi-C libraries. (**B**) Correlations between the Hi-C data generated in this study and previously published (*48, 49*). Correlations of contact matrices are determined by Pearson’s correlation coefficient (*R*). (**C**) Hi-C interaction frequency heatmaps at 500 kb resolution for WT (upper) and *p35S::H2B.8-eGFP* (lower) seedlings. (**D**) Intrachromosomal interaction probability against genomic distance for WT (blue) and *p35S::H2B.8-eGFP* (red) Hi-C libraries.

**Fig. S7.**
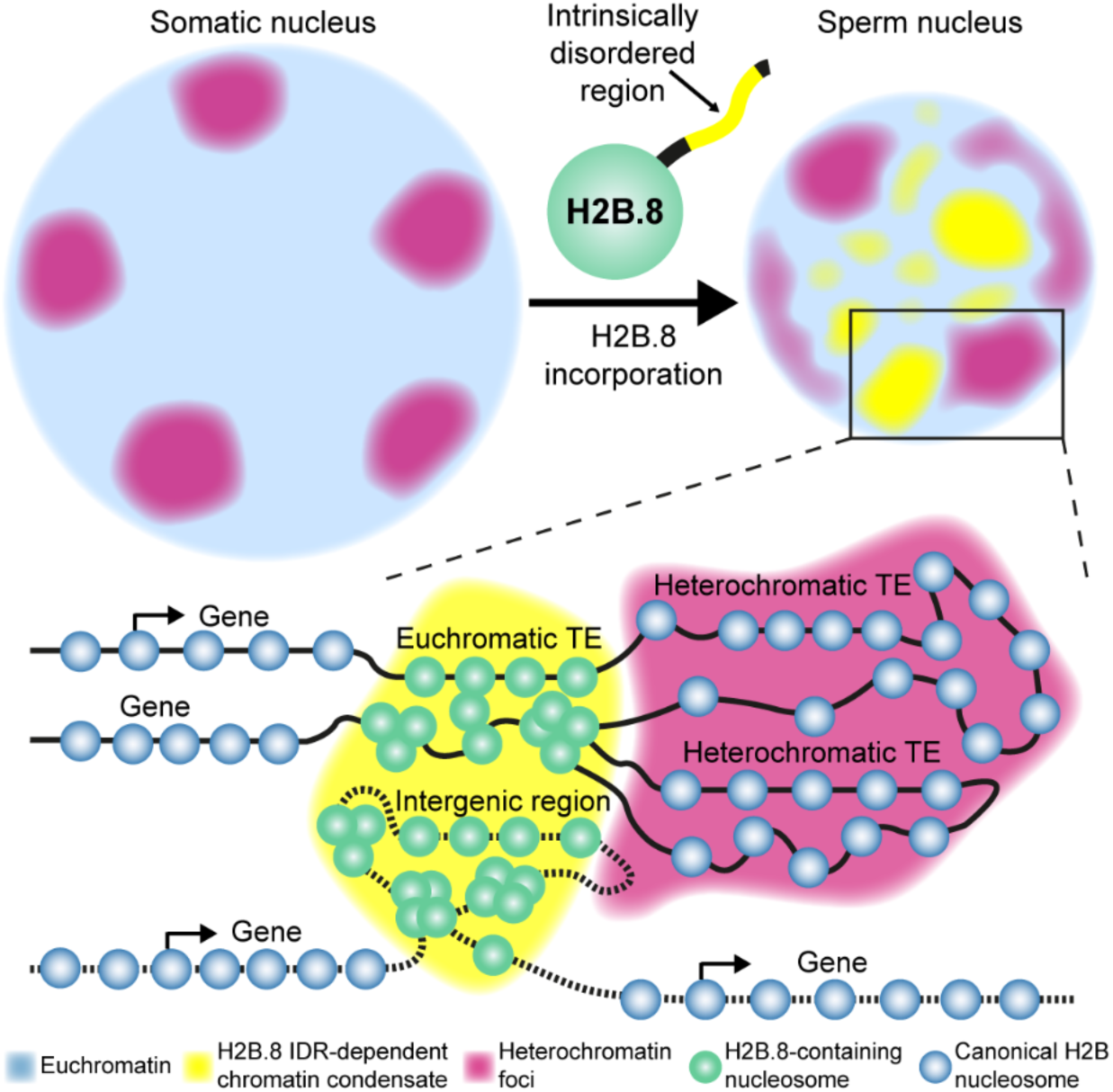
Mechanism of H2B.8-mediated sperm condensation. H2B.8 drives sperm nuclear compaction via the formation of chromatin condensates (yellow), which is dependent on an intrinsically disordered domain (IDR) of H2B.8 conserved among flowering plants. Unlike typical chromatin condensation mechanisms, H2B.8-induced condensation does not inhibit transcription. Condensation is achieved by the specific deposition of H2B.8 into inactive AT-rich chromatin, which alters higher-order chromatin architecture to effectively compact the nucleus without sacrificing transcription. H2B.8-mediated chromatin aggregation disperses heterochromatin foci (pink), suggesting interactions between the euchromatic (yellow) and heterochromatic (pink) chromatin condensates in the nucleus.

**Table S1. List of H2B.8 homologs identified in flowering plant species.**

File attached.

**Table S2.**
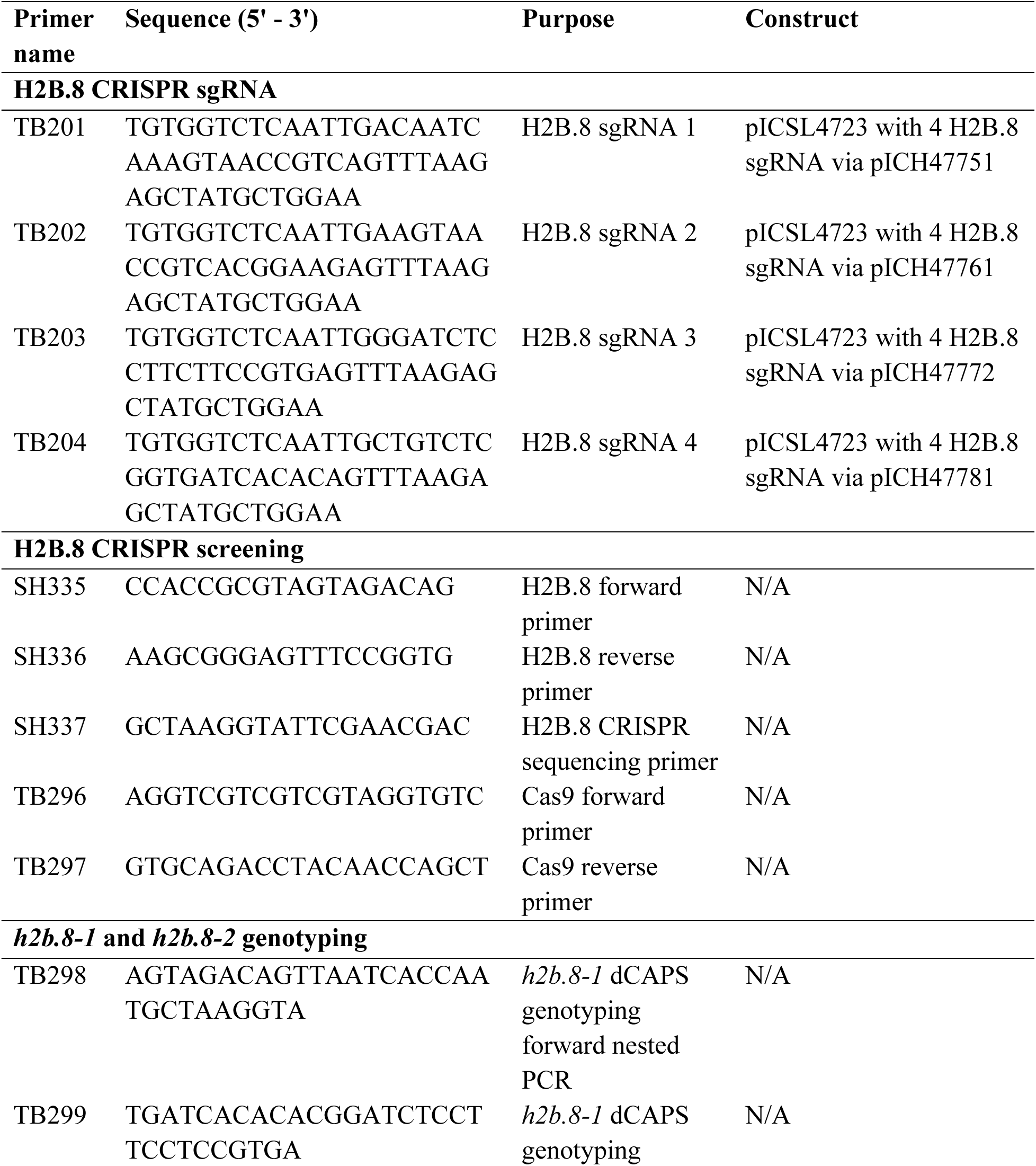
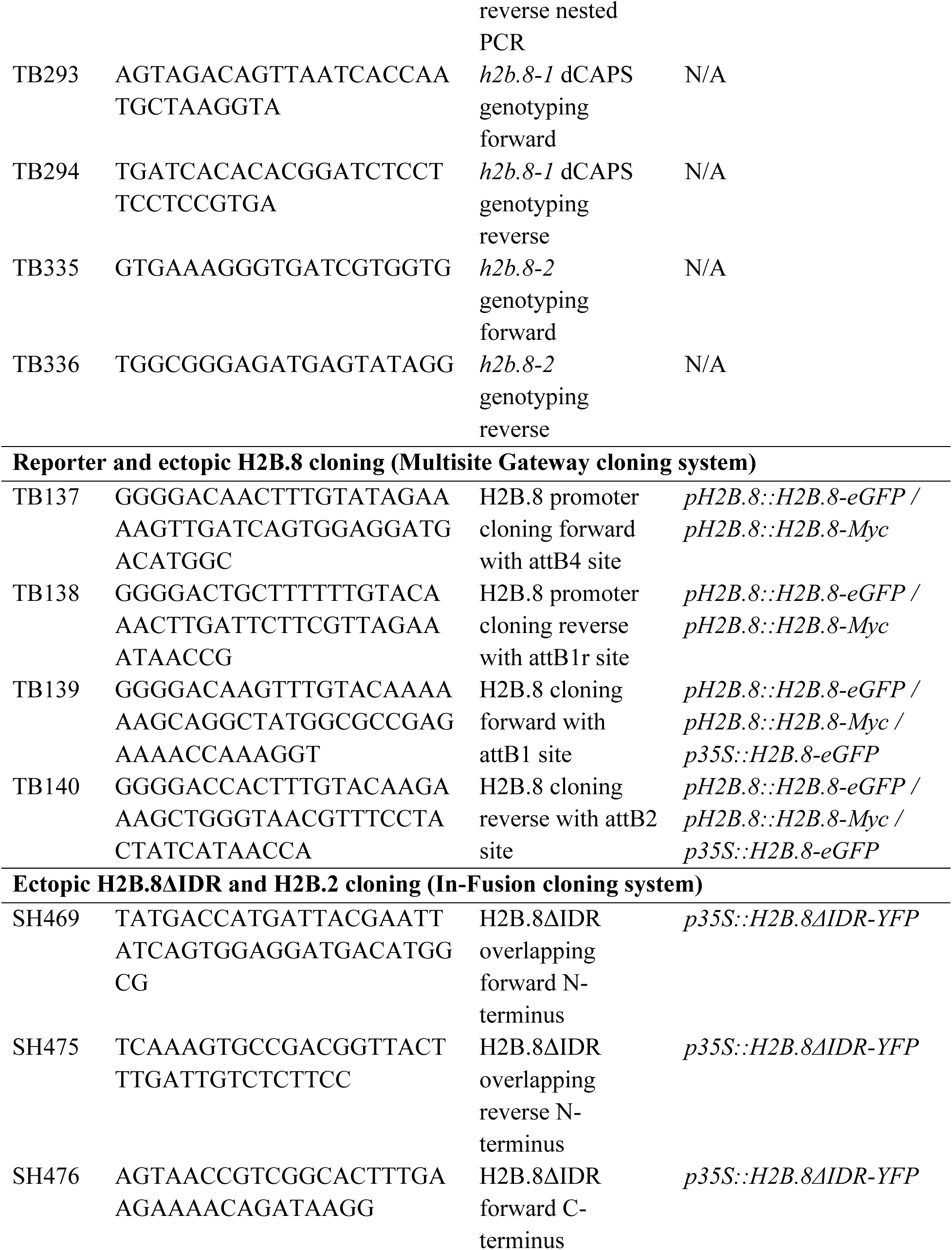
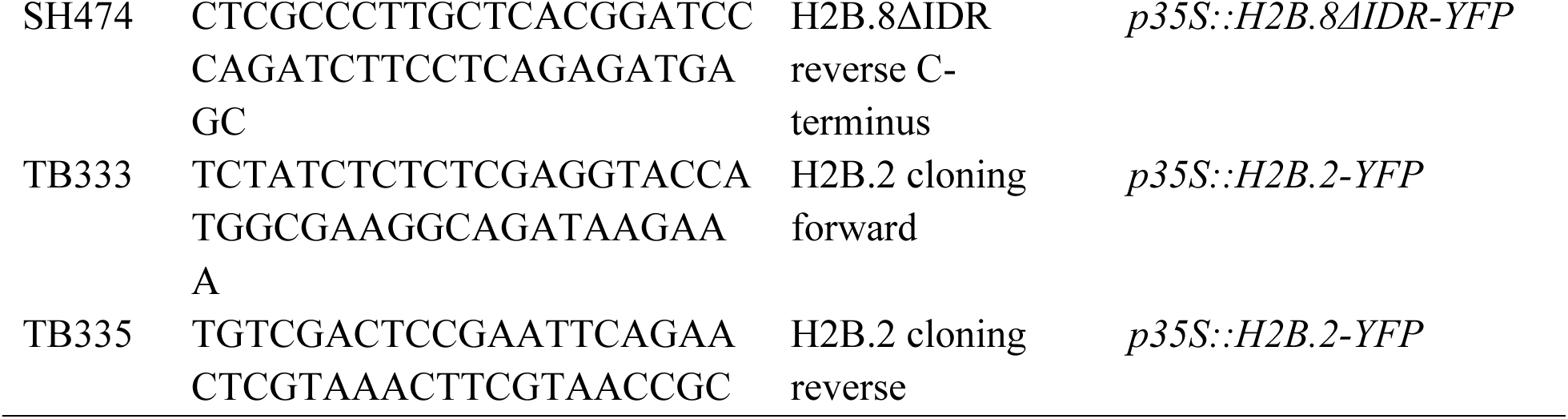
List of primers used in this study.

